# Decreased myelin-related gene expression in the nucleus accumbens during spontaneous neonatal opioid withdrawal in the absence of long-term behavioral effects in adult outbred CFW mice

**DOI:** 10.1101/2023.08.04.552033

**Authors:** Kristyn N. Borrelli, Kelly K. Wingfield, Emily J. Yao, Catalina A. Zamorano, Katherine D. Sena, Jacob A. Beierle, Michelle A. Roos, Huiping Zhang, Elisha M. Wachman, Camron D. Bryant

**Affiliations:** Laboratory of Addiction Genetics, Department of Pharmacology, Physiology & Biophysics, Boston University Chobanian and Avedisian School of Medicine, 72 E. Concord St., L-606B, Boston, MA 02118; T32 Biomolecular Pharmacology PhD Program, Boston University Chobanian and Avedisian School of Medicine; Graduate Program for Neuroscience, Boston University, 610 Commonwealth Av, Boston, MA 02215; Boston University’s Transformative Training Program in Addiction Science, Boston University Chobanian & Avedisian School of Medicine, 72 E. Concord St., L-317, Boston, MA 02118; Boston University’s Undergraduate Research Opportunity Program, George Sherman Union, 775 Commonwealth Av, 5th floor, Boston, MA 02215; Department of Psychiatry, Boston University Chobanian and Avedisian School of Medicine, 72 E. Concord St., Boston, MA 02118; Department of Pediatrics, Boston University Chobanian and Avedisian School of Medicine and Boston Medical Center, 1 Boston Medical Center Pl, Boston, MA 02118

**Keywords:** opiate, neonatal abstinence syndrome, neonatal opioid withdrawal syndrome, swiss webster, RNA-seq, opioid use disorder, ICSS

## Abstract

Prenatal opioid exposure is a major health concern in the United States, with the incidence of neonatal opioid withdrawal syndrome (NOWS) escalating in recent years. NOWS occurs upon cessation of *in utero* opioid exposure and is characterized by increased irritability, disrupted sleep patterns, high-pitched crying, and dysregulated feeding. The main pharmacological strategy for alleviating symptoms is treatment with replacement opioids. The neural mechanisms mediating NOWS and the long-term neurobehavioral effects are poorly understood. We used a third trimester-approximate model in which neonatal outbred pups (Carworth Farms White; CFW) were administered once-daily morphine (15 mg/kg, s.c.) from postnatal day (P) day 1 through P14 and were then assessed for behavioral and transcriptomic adaptations within the nucleus accumbens (NAc) on P15. We also investigated the long-term effects of perinatal morphine exposure on adult learning and reward sensitivity. We observed significant weight deficits, spontaneous thermal hyperalgesia, and altered ultrasonic vocalization (USV) profiles following repeated morphine and during spontaneous withdrawal. Transcriptome analysis of NAc from opioid-withdrawn P15 neonates via bulk mRNA sequencing identified an enrichment profile consistent with downregulation of myelin-associated transcripts. Despite the neonatal behavioral and molecular effects, there were no significant long-term effects of perinatal morphine exposure on adult spatial memory function in the Barnes Maze, emotional learning in fear conditioning, or in baseline or methamphetamine-potentiated reward sensitivity as measured via intracranial self-stimulation. Thus, the once daily third trimester-approximate exposure regimen, while inducing NOWS model traits and significant transcriptomic effects in neonates, had no significant long-term effects on adult behaviors.

**HIGHLIGHTS:**

- We replicated some NOWS model traits via 1x-daily morphine (P1-P14).
- We found a downregulation of myelination genes in nucleus accumbens on P15.
- There were no effects on learning/memory or reward sensitivity in adults.

## 1. Introduction

Prenatal exposure to opioids and subsequent consequences, including neonatal opioid withdrawal syndrome (**NOWS**) and long-term developmental effects represents a growing public health concern in the United States. In 2020, the Healthcare Cost and Utilization Project (HCUP) reported that approximately more than 59 newborns are diagnosed with NOWS every day (HCUP Fast Stats, 2021).

Clinical cases of NOWS following *in utero* opioid exposure are characterized by symptoms reflecting disruptions to the central nervous system (e.g., high-pitched crying, fragmented sleep, high muscle tone, and tremors), autonomic nervous system (e.g., sweating, increased body temperature, yawning, sneezing, nasal congestion, and heightened respiratory rate), and gastrointestinal tract (trouble feeding, vomiting, and loose stools) (Patrick et al. 2020). First-line pharmacological treatment of NOWS is typically with an opioid medication such as mu opioid receptor agonists (e.g., morphine, methadone, or buprenorphine), and second-line pharmacologic agents including phenobarbital and clonidine (Hall et al. 2018; Kraft et al. 2017; Siu and Robinson 2014; Wachman et al. 2011; Wachman, Schiff, and Silverstein 2018; Wachman and Werler 2019). Despite the availability of current treatment options, NOWS is a significant financial burden, as hospital costs are nearly 7-times greater than other newborn stays (HCUP Fast Stats, 2021; Winkelman et al. 2018).

The potential for additional non-opioid pharmacological treatments could reduce the time to recovery from NOWS and thus shorten hospital stays and associated costs. Furthermore, the long-term effects of opioid exposure on neurodevelopment and behavior remain poorly understood (McPhail et al. 2021). Multiple studies have reported adverse effects of perinatal opioid exposure on cognitive function, motor development and unfavorable behavior observed in toddlers and children under 12. (Bakhireva et al. 2019; C. R. Bauer et al. 2020; Bernstein et al. 1984; Johnson and Rosen 1982; Konijnenberg, Sarfi, and Melinder 2016; Wouldes and Woodward 2020). Thus, additional basic and clinical research are needed to inform mechanisms underlying NOWS and developmental perturbations to improve treatments and long-term outcomes.

As there does not appear to be a correlation between maternal opioid dose and NOWS severity, genetic and epigenetic factors are suspected to contribute to the significant variability in neonatal outcomes (Wachman et al. 2013; Wachman, Schiff, and Silverstein 2018; Cole, Wegner, and Davis 2017; Brogly et al. 2014). Genetic studies in both mothers and neonates exposed to opioids in utero revealed single-nucleotide polymorphisms in genes related to addiction, opioid metabolism, and stress, and were associated with differences in treatment outcomes (Borrelli et al. 2022; Cole, Wegner, and Davis 2017; Lewis, J, and Js 2015; Wachman et al. 2013; Wachman, Schiff, and Silverstein 2018). Clinical variables such as gestational age, biological sex, breastfeeding, rooming-in with the parent, type of maternal opioid exposure and maternal polysubstance use are also associated with differences in NOWS occurrence and severity, and treatment strategy (Desai et al. 2015; Haight 2018; Jones et al. 2013; Wachman et al. 2011).

Given the limitations of environmental variability and the difficulty in implementing longitudinal study designs in humans, rodent models of perinatal opioid exposure offer an efficient means to study the acute, neurobehavioral adaptations that manifest during perinatal opioid exposure/withdrawal and the long-term effects in adolescence and adulthood. In a recent report where we employed a mouse model for NOWS adapted from a previous third trimester-approximate protocol (Robinson S et al., 2020; Genes, Brain and Behavior), we identified sex-specific transcriptional adaptations in the brainstem following postnatal exposure to morphine (Borrelli, Yao, et al. 2021).

The first two postnatal weeks of opioid exposure in rodents serve as a model for several similar neurodevelopmental processes that occur in human fetuses during the third trimester (Semple et al. 2013) and permit precise control over the amount of opioid exposure across individual pups. Furthermore, the third trimester is a critical exposure period for presentation of NOWS in humans (Desai et al. 2015). Thus, in this study, we employed a third trimester-approximate exposure model with once-daily morphine (15 mg/kg, s.c.) from postnatal day (P) 1 through P14 in outbred Swiss Webster Carworth Farms White (CFW) mice. Our purpose was multi-fold. First, we switched from a twice-daily regimen (Borrelli et al., 2021) to a once-daily regimen to improve survival rate and determine if this would be sufficient to induces NOWS traits. Second, we wanted to examine transcriptomic adaptations within the NAc during the expression of NOWS model symptoms as previous analysis was limited to the brainstem. The NAc is a critical brain region implicated in the negative affective state during spontaneous opioid withdrawal (Welsch et al. 2020). Third, we wanted to examine the long-term consequences of daily, third trimester-approximate exposure in adulthood on spatial memory, reward sensitivity, and cue-conditioned fear learning. The results indicate a more subtle set of NOWS model traits during spontaneous morphine withdrawal following once-daily, third trimester-approximate morphine exposure and a unique transcriptomic signature showing a downregulation of genes involved in myelination. Furthermore, there was no long-term impact of opioid exposure on learning and reward sensitivity in young adult mice.

## 2. Materials and methods

### 2.1. Mice

All experiments in mice were conducted in accordance with the NIH Guidelines for the Use of Laboratory Animals and were approved by the Institutional Animal Care and Use Committee at Boston University. Outbred Carworth Farms White (CFW) mice (Swiss Webster) were purchased from Charles River Laboratory at 6 weeks of age. Breeders were paired after two weeks of habituation to the vivarium. Each breeder mouse was from a different litter in order to minimize relatedness within breeder pairs (Parker et al. 2016). Laboratory chow (Teklad 18% Protein Diet, Envigo, Indianapolis, IN, USA) and tap water were provided *ad libitum.* Breeder cages were provided with nestlets. A maximum of two litters per breeder pair were used in the study. Mice were maintained on a 12:12 light-dark cycle (lights on, 0630 h). Phenotyping was conducted in both female and male mice during the light phase between 0900 h and 1200 h. All behavioral experiments were performed by female experimenters, thus controlling for variance attributable to experimenter sex (Bohlen et al. 2014; Sorge et al. 2014).

### 2.2. Morphine treatment regimen from P1 through P14

Morphine sulfate pentahydrate (Sigma-Aldrich, St. Louis, MO USA) was dissolved in sterilized physiological saline (0.9%) in a 0.75 mg/ml morphine sulfate pentahydrate solution for systemic administration via subcutaneous (s.c.) injections (15 mg/kg, 20 ul/g volume). For P1 through P14, injections were administered once daily at 1700 h. We used a split-litter design wherein one-half of the pups were randomly assigned to saline (20 ul/g, s.c.) and the other half randomly assigned to morphine (15 mg/kg, s.c.) within the same litter. Having both drug treatment groups within each litter (MOR and SAL-treated) helps mitigate potentially confounding, between-treatment variance caused by relatedness, maternal care, and the overall within-cage environment. Behavioral phenotyping was performed on P7 and P14 16 h after the previous morphine administration, allowing for the emergence of spontaneous opioid withdrawal (Boasen et al. 2008). Separate cohorts of mice were used for transcriptome analysis (RNA-seq), Barnes maze/fear conditioning, and intracranial self-stimulation (ICSS). Body weight, nociception, and ultrasonic vocalizations (USV) recordings were collected from all cohorts. Cohorts used for adult phenotyping were weaned at P20 and housed with same-sex littermates until P60.

### 2.3. General Procedures: P7 and P14 Phenotyping

Prior to testing, pups were transferred to a holding cage placed on top of an electric heating pad. Dams and sires remained in their home cages in the testing room. Activity in sound attenuating chambers during ultrasonic vocalization recording and on the hot plate were video-recorded using infrared cameras (Swann Communications U.S.A. Inc., Santa Fe Springs, CA, USA) and later tracked with ANY-maze V tracking software (Stoelting Co., Wood Dale, IL, USA).

### 2.4. Ultrasonic Vocalizations (USVs) in P7 and P14 pups

Ultrasonic vocalizations were recorded as previously described (Borrelli, Yao, et al. 2021). On P7 and P14, pups were placed into Plexiglas arenas (43 cm L x 20 cm W x 45 cm H; Lafayette Instruments, Lafayette, IN, USA) within sound-attenuating chambers (Med Associates, St. Albans, VT, USA). USVs were recorded using Ultrasound Recording Interface (Avisoft Bioacoustics UltrasoundGate 816H, Glienicke, Germany). USVs were recorded for 10 min on P7 and 15 min on P14. We employed a shorter time frame on P7, due to the reduced ability to regulate body temperature and a longer time frame on P14 because of the pups increased ability to regulate body temperature and thus the opportunity to collect additional data. Following recordings, pups were returned to the holding cage with a heating pad prior to nociceptive testing.

### 2.5. Locomotor activity during USV recordings in P7 and P14 pups

Video recordings were captured from Swann video cameras mounted on the ceiling of the sound attenuating chambers, above the Plexiglas arenas (Borrelli et al., 2021). We used AnyMaze video tracking software to quantify distance traveled (m) over the 10-min (P7) or 15-min (P14) intervals.

### 2.6. Thermal nociception in P7 and P14 pups

Thermal nociception was determined as described (Borrelli, Yao, et al. 2021). Pups were removed from the holding cage and placed in a Plexiglas cylinder (15 cm diameter; 33 cm high) on a 52.5°C hot plate (IITC Life Science INC., Woodland Hills, CA, USA). On P7, the nociceptive response was the latency for the pup to roll onto its back as an avoidance response to thermal exposure of the paws (they are unable to lick at that age). The nociceptive response on P14 was the latency to lick the forepaw or hindpaw or jump. Pups were removed from the hotplate immediately following a pain response or after a cut-off time of 30 s (P7) or 60 s (P14). Following hot plate testing, pups were gently scruffed and the distal half of the tail was rapidly submerged in a 48.5°C hot water bath. The latency to flick the tail was used as the nociceptive response.

### 2.7. Tattooing in P7 mice

Following thermal nociception testing on P7, pups were tattooed on their tails for identification purposes (ATS-3 General Rodent Tattoo System, AIMS™, Hornell, NY, USA). Pups were then injected with morphine or saline and returned to their home cage.

### 2.8. Statistical analysis of behavior

Data analysis was performed in R (https://www.r-project.org/) and IBM SPSS Statistics 27. Body weight, hot plate, tail withdrawal, hot plate velocity, USVs per min, time freezing per min, total time freezing, and ICSS responses/rewards were analyzed using multifactorial repeated measures (RM) ANOVAs, with Morphine Treatment and Sex as between-subjects factors and Age/Day, Time, or Drug Dose (for ICSS) as repeated measures.

Data residuals for each phenotype were assessed for normality prior to parametric analysis. In the case of a non-normal distribution of the residuals (Shapiro-Wilk test for normality, p<0.05), data were percentile rank-normalized (Templeton 2011) and the Shapiro-Wilk test was repeated to confirm successful normalization (p>0.05). To facilitate interpretation of the data, normalized data are presented as raw measurement values instead of normalized values. Mauchly’s test of sphericity was performed for repeated measures with more than two levels (body weight and USV analysis). If the assumption of sphericity was violated, a Greenhouse-Geisser correction was implemented. Homogeneity of variances was confirmed using Levene’s Test of Equality of Error Variances. Post-hoc comparisons were performed in the case of a significant interaction to determine group differences. Either pairwise comparisons with Bonferroni correction or Tukey’s Honestly Significant Difference (HSD) tests were conducted for multiple comparisons.

### 2.9. Bulk mRNA sequencing (RNA-seq) and differential gene expression analysis of nucleus accumbens tissue

On P15, at approximately 16 h after the final morphine injection (during a state of spontaneous morphine withdrawal), pups were sacrificed, and brains were quickly removed for tissue collection. Punches (2 mm thick) from the NAc containing both core and shell were transferred to tubes containing RNA-later, stored at 4°C for 3 days, and then transferred to new tubes to store at -80°C until RNA extraction. RNA was extracted using Trizol (Qiagen), ethanol precipitation, filtering columns (Qiagen), and elution with sterile, double deionized water (Yazdani et al. 2015). RNA library preparation (poly-A selection) and RNA-seq were performed in two batches. The first batch (8 samples: 4 SAL; 4 MOR) was processed at the Boston University Microarray and Sequencing Resource (BUMSR) Core Facility on an Illumina NextSeq2000 PE-100 (paired-end 100-bp) flow cell. The second batch (28 samples: 14 SAL; 14 MOR) was processed at the University of Chicago Genomics Facility on an Illumina NovaSEQ6000 using a PE-100 flow cell. Reads from both batches were trimmed for quality using Trimmomatic (Bolger, Lohse, and Usadel 2014). Trimmed reads were then aligned to the mm10 mouse reference genome (Ensembl) to generate BAM files for alignment using STAR (Dobin et al. 2013). The featureCounts read summarization program was used to count reads mapping to the “exon” feature in a GTF file obtained from Ensembl (GRCm38). The average library size was 49 million reads (ranging from 42-82 million). Genes with counts per million (cpm) < 0.5 in at least half of the samples were removed prior to DEG analysis. Read counts were normalized using the ‘voom’ function in the limma R package (v.3.46.0) (Ritchie et al. 2015). Differentially expressed genes (DEGs) were determined using a linear model with ‘Opioid Treatment’, ‘Sex’, ‘Batch’, and ‘Litter’ as covariates. The design matrix was set up as the following: (∼ Opioid Treatment + Sex + Batch + Litter). The effect of ‘Opioid Treatment’ was isolated to identify DEGs. DEGs were visualized with volcano plots (Blighe, K, S Rana, and M Lewis 2018).

### 2.10. Enrichment and gene network analysis

Downregulated genes with an unadjusted p-value < 0.01 and log_2_FC < -0.26 were included in pathway enrichment and gene network analysis within the gene ontology (GO) biological process, molecular function, and cellular component terms. Cytoscape software (https://www.cytoscape.org) was used to perform pathway enrichment analysis of downregulated DEGs, and to generate and plot known gene interaction pathways using the STRING Mus musculus database.

### 2.11. Barnes maze

Adult mice (P96 – P100) were trained on an abbreviated Barnes maze protocol (Attar et al. 2013). The apparatus consisted of a circular, wall-less, and elevated platform that is 95 cm tall and 92 cm in diameter with 20 holes (5 cm diameter) equally spaced around the perimeter (2 cm from edge) (MazeEngineers, Skokie, IL). Four visual cues of different color were placed 30 cm from the edge of the maze and were spaced evenly around the maze perimeter. All mice from the same cage (2-5 mice per cage; n = 56) were placed in holding cages during habituation and training trials. Habituation was performed for all mice/cages prior to training trials. Between each trial, the maze and escape cage were thoroughly cleaned with 70% ethanol and the maze was randomly rotated to remove odor cues. The location of the escape hole was changed after every two cages. On Day 1, mice first underwent a habituation trial, during which they were placed in the center on the maze under a clear Plexiglas cylinder (15 cm diameter; 33 cm high) for 30 s and were then guided to the target hole with an escape cage underneath. Mice were allotted 1 min to enter the escape cage independently before they were gently guided into it. The trial ended after the mouse spent 1 min in the escape cage. Two training trials were performed following habituation on training day 1 and three training trials were performed 24 h later on training Day 2. During training trials, mice were placed under an opaque cylinder in the center of the maze for 15 s. The cylinder was lifted, and mice were allowed to explore the maze for 2 min. If a mouse entered the escape cage, it was allowed 1 min in the cage before the end of that training trial. If a mouse did not enter the escape cage within 2 min, it was guided to the escape hole under the clear cylinder and allowed 1 min to enter, after which it was gently nudged into the escape cage. Again, all mice were allowed 1 min in the escape cage after entering. Training trials on Day 1 and Day 2 were performed one cage at a time, such that all trials were completed for mice within one cage before proceeding to the next. Forty-eight h after the second training day, probe trials were performed. The escape hatch was removed for probe day testing. During probe trials, mice were placed in the center of the maze under an opaque cylinder for 15 s, then the cylinder was lifted and the mice were given 2 min to freely explore the maze. Videos were recorded of all training and probe trials and were later analyzed via AnyMaze software to determine the primary latency for the mouse to investigate the target hole, the time spent in the target quadrant, and the primary holes searched prior to identifying the target hole.

### 2.12. Fear conditioning

Approximately 1 week after completing Barnes maze testing, mice underwent fear conditioning training within operant boxes (15.24 × 13.34 × 12.7 cm; ENV-307A-CT; Med Associates) inside closed sound-attenuating cubicles (ENV-022V; Med Associates). Day 1 of training consisted of 3 min habituation to the conditioning context followed by the presentation of a 20 s auditory tone (75 dB, 2 kHz) (ENV-323HAM; Med Associates). At the end of the tone, a 2 s 0.5 mA foot-shock (ENV-414; Med Associates) was delivered. Each foot-shock was followed by a 3 min interval with no auditory cue or foot-shocks. A total of 3 tone/foot-shock pairings were delivered. On day 2, mice were placed in the same context as day 1 without any auditory cues or foot-shocks. To strengthen the experience of a novel environment, on day 3, mice were placed in the same chambers as on day 1 and 2 with altered floor textures and wall color. Peppermint oil was used an odor cue to further solidify the new testing context. Mice underwent a 3 min context-habituation period followed by the same tone delivered on day 1. No foot-shocks were delivered on day 3. A total of 3 tones were presented, each followed by a 3 min interval with no tone.

### 2.13. Intracranial self-stimulation (ICSS)

A separate cohort of adult mice underwent ICSS testing to assess sensitivity to reward stimulation and reward potentiation in the presence of increasing doses of methamphetamine (Borrelli et al., 2021), a potent dopamine releaser within the mesolimbic reward pathway (C. T. Bauer et al. 2013). Surgeries, training, and testing were performed as described (Borrelli, Langan, et al. 2021), using a previously published protocol (Carlezon and Chartoff 2007). Briefly, mice underwent stereotaxic surgery to implant a bipolar stimulating electrode (Plastics1, Roanoke, VA; MS308) to the medial forebrain bundle. Mice were trained on a fixed-ratio 1 (FR1) schedule to respond for stimulation via a wheel manipulandum (ENV-113AM; MedAssociates, St. Albans, VT, USA) within an operant chamber (15.24 × 13.34 × 12.7 cm; ENV-307A-CT; MedAssociates). For every ¼ turn of the response wheel, a 500-ms train of square-wave, cathodal current (100-ms pulses) was delivered at a constant frequency of 142 Hz through a stimulator (PHM-150B/2; MedAssociates). Each stimulation was followed by a 500-ms time-out period during which responses were counted but not reinforced by stimulation. The current intensity was adjusted for each subject to the lowest value that produced > 500 responses during a 45-min period across at least three consecutive training sessions, and the minimum effective current for each subject was held constant throughout the rest of the study. All behavioral procedures were performed using Med-PC V software (MedAssociates).

Mice were then introduced to stimulation that was delivered over a series of 15 descending frequencies (log0.05 steps, 142–28 Hz). Each frequency trial consisted of 5 s of non-contingent priming stimulation, a 50-s response period during which responses were rewarded with 500-ms stimulation, and a 5-s time-out period. Each response-contingent stimulation was followed by a 500-ms timeout. This sequence was repeated for all 15 frequencies, and the procedure was repeated two more times in the same session. The number of responses (including those emitted during timeout periods) and stimulations delivered were recorded for each trial. Training was repeated until mice consistently responded at maximum rates during the highest frequency trials and did not respond during the lowest frequency trials. Subjects were required to demonstrate less than 15% between-day variability in mean response threshold for at least three consecutive training days prior to testing with methamphetamine (MA).

Testing was performed every other day and all mice received the same escalating doses of MA on these days using a within-subjects design. Mice were first assessed for baseline, drug-naïve ICSS responding via a rate-frequency procedure that was identical to what was used during training (see above). Upon completion of the 45-min baseline procedure, mice were removed from the operant chambers, injected with MA (0.25, 0.50, 1.0, 2.0 mg/kg; i.p.), and placed individually into clean cages with fresh bedding. Ten min post-MA injection, mice were placed back into the operant chambers and underwent the same protocol used during baseline assessment (3 passes of 15 min).

## 3. Results

Sample sizes for each experiment are included in the figure panels and/or legends.

### 3.1. Body weight

A schematic of the treatment assignment and timeline for phenotyping is provided in **Figure 1A**. Reduced body weight is a classic manifestation of infants born with NOWS. Once daily injections of morphine (15 mg/kg, s.c.) or saline (20 ul/g, sc.) from P1-P14 resulted in reduced body weight gain from P2-P14 (**Figure 1B**). Body weight data were normalized prior to analysis (resulting Shapiro-Wilk statistic *p* > 0.05) as Mauchly’s test indicated the sphericity assumption had been violated (χ^2^_90_ = 3850.49; *p* < 0.001) so the Greenhouse-Geisser correction was applied (ε = 0.151). There were no significant differences in body weight on P1 prior to the first injections. Sex did not interact with morphine-induced weight deficits. There were main effects of Drug (*F*_1,210_ = 160.13; \*\*\**p* < 0.001), Sex (*F*_1,210_ = 5.3; *p* = 0.02), and Day (*F*_1.96,412.22_ = 7293.67; \*\*\**p* < 0.001) as well as Drug x Day (*F*_1.96,412.22_ = 175.28; \*\*\**p* < 0.001) and Sex x Day (*F*_1.96,412.22_ = 4.79; *p* = 0.009) interactions. Post-hoc multiple comparison tests with Bonferroni correction were performed to compare effect of Treatment (saline vs morphine pups; sex-collapsed) and the effect of Sex (males vs. females; treatment-collapsed) on each postnatal day. For all days except P1 (*p* = 1; prior to any drug treatment), morphine pups had significantly lower body weights than saline pups (P2: \*\*\**p* = 0.0002; P3-P14: \*\*\*\**p* < 0.0001; **Figure 1B**). There were no significant differences between females and males (all *p* = 1.0).

**Figure 1.**
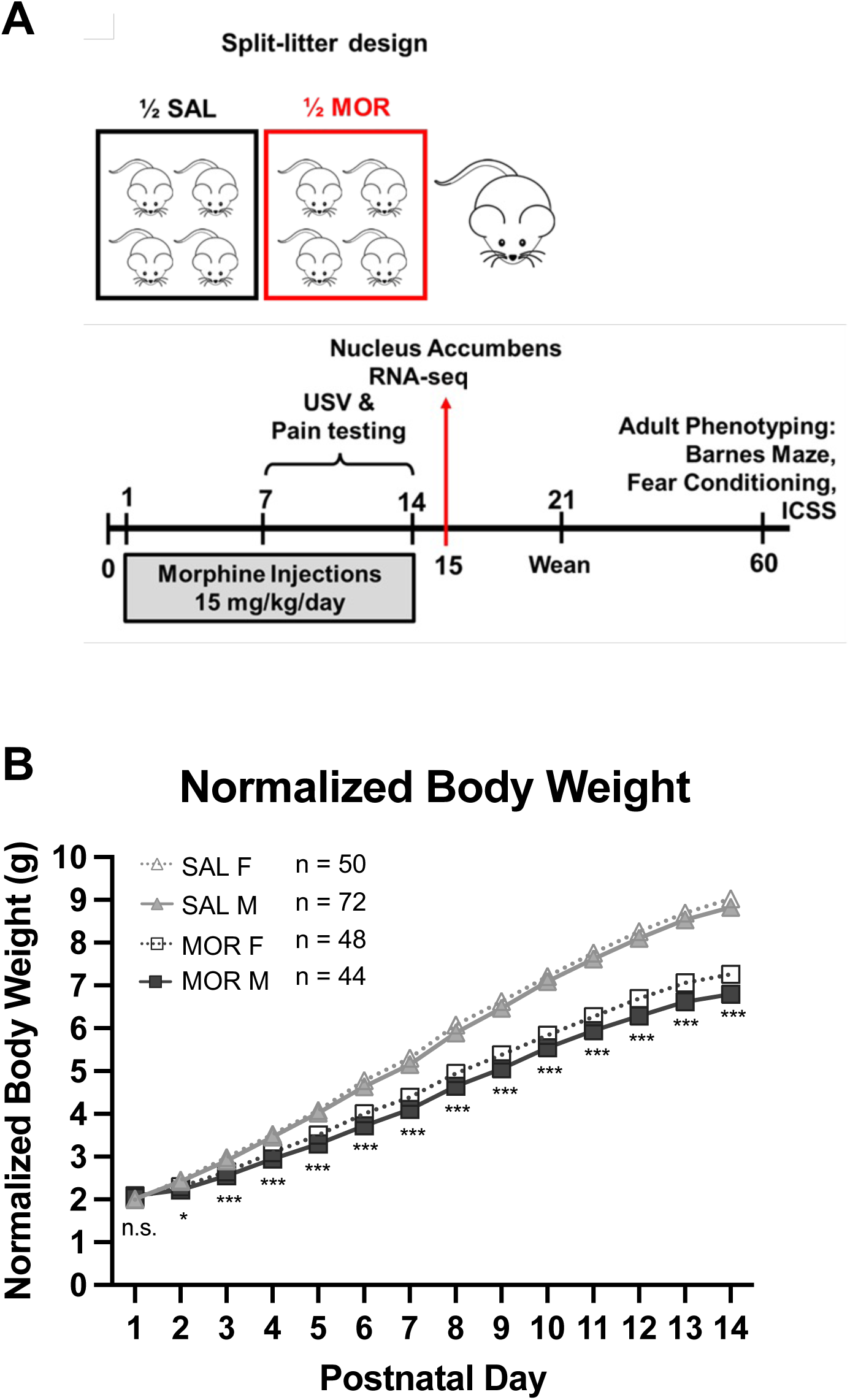
Reduced body weight from P2 through P14 following once daily morphine exposure (15 mg/kg; s.c.). **A.** Schematic representing the split-litter design (top) and timeline of morphine administration, tissue collection, and behavioral phenotyping. **B.** Asterisks indicate differences between morphine and saline groups (sex-collapsed; n.s. = not significant, *p* < 0.01 = *, *p* < 0.001 = ***). All *p*-values reflect Bonferroni adjustment for multiple comparisons across days. Data are plotted as the mean ± SEM. Sample sizes by Treatment and Sex are listed next to the figure legend.

### 3.2. Thermal nociception

All nociception data, including hot late latency and velocity and tail withdrawal latency were normalized prior to analysis (Shapiro-Wilk *p* values > 0.05). For the hot plate, morphine pups displayed thermal hyperalgesia as evidenced by a shorter latency to elicit a nociceptive response (**Figure 2A**). Because there was no effect of prior morphine treatment on hot plate velocity measured on P7 and P14 (**Figure 2B**), this indicated that the hot plate nociceptive response was not confounded by concomitant locomotor activity on the hot plate. Specifically, morphine-treated pups exhibited robust thermal hyperalgesia on P7 and P14. There were main effects of Day (*F*_(1,211)_ = 252.27, \*\*\**p* < 0.001) and Morphine Treatment (*F*_(1,211)_ = 65.74, \*\*\**p* < 0.001). There was no effect of Sex (*F*_(1,211)_ = 1.72, *p* = 0.19) and no significant interactions (all *p* > 0.07; Sex x Day: *F*_(1,211)_ = 3.30, *p* = 0.07; **Figure 2A**). Tukey’s post-hoc indicated a significant reduction in hot plate latencies in morphine treated pups on P7 and P14 (\**p* < 0.05). For locomotor velocities (cm/s) on the hot plate prior to nociceptive responses, there was a main effect of Day (*F*_(1,99)_ = 7.193, \*\*\**p* < 0.001) but no effects of Morphine Treatment (*F*_(1,99)_ = 0.203, *p* = 0.65) or Sex (*F*_(1,99)_ = 0.703, *p* = 0.40) and no interactions (all *p* ≥0.24).

**Figure 2:**
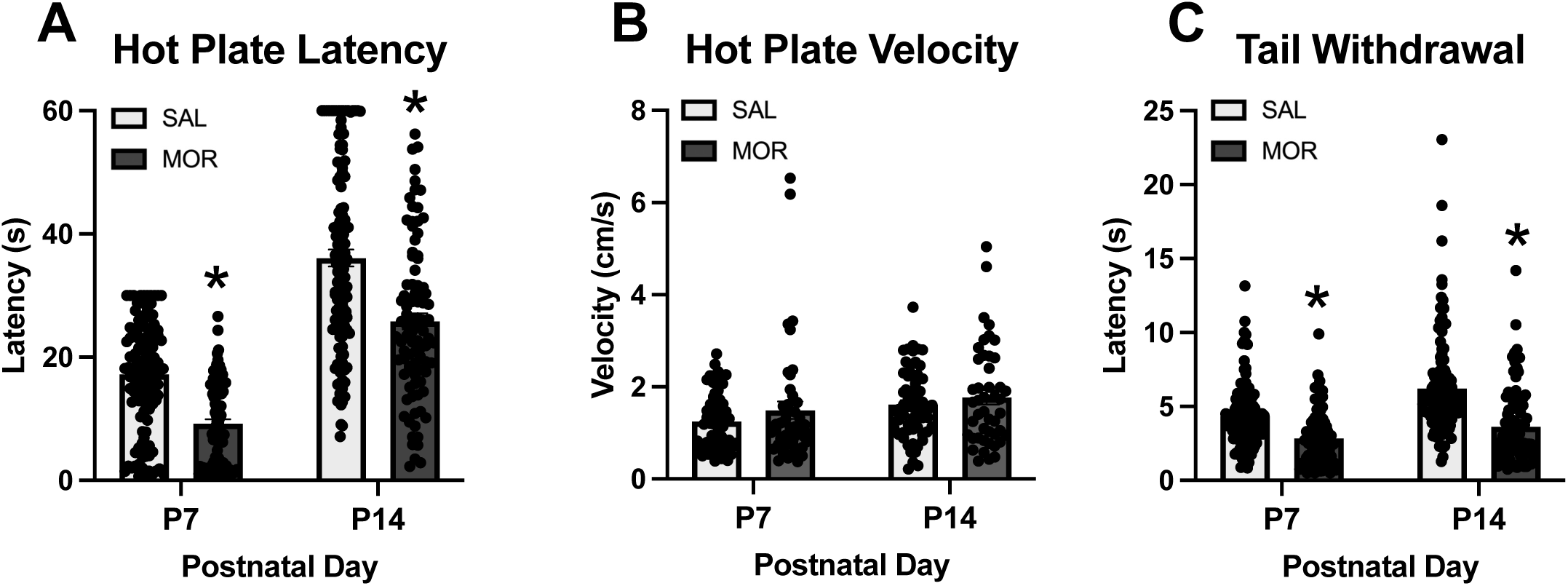
Thermal hyperalgesia in the hot plate and tail withdrawal assays during spontaneous withdrawal on P7 and P14 following once daily morphine exposure (15 mg/kg, s.c.) from P1-P14. Data are plotted as the mean ± SEM. **A.** Hot plate latencies: Main effects of Day (\*\*\**p* < 0.001) and Morphine Treatment (\*\*\**p* < 0.001), but no interactions. Morphine treated pups showed decreased latencies on P7 and P14 (*Tukey’s p < 0.05). P7 SAL: n = 123 (51 females, 72 males); SAL P14: n = 123 (51 females, 72 males); MOR P7: n = 92 (48 females, 44 males); MOR P14, n = 92 (48 females, 44 males). **B.** Locomotor velocities (cm/s) on the hot plate prior to nociceptive responses: No effect of Morphine Treatment but an effect of Day (*p* < 0.001). SAL P7 – P14: n = 60 (24 females, 36); MOR P7 – P14: n = 46 (27 females, 19 males). **C.** Tail withdrawal latencies: Effect of Morphine Treatment (\*\*\**p* < 0.001), Day (\*\*\**p* < 0.001), and interaction (\*\**p* = 0.004). Reduced latencies were observed in morphine treated pups on both P7 and P14 (*Tukeys; p < 0.05). SAL P7: n = 123 (51 females, 72 males); SAL P14: n = 119 (49 females, 70 males); MOR P7 – P14: n = 92 (48 females, 44 males).

For tail withdrawal, morphine pups also displayed thermal hyperalgesia (shorter tail flick latency; **Figure 2C**). Specifically, there were main effects of Day (*F*_(1,205)_ = 33.64,\*\*\**p* < 0.001) and Morphine Treatment (*F*_(1,205)_ = 76.53, \*\*\**p* < 0.001), and a Morphine Treatment x Day interaction *F*_(1,205)_ = 8.26, \*\**p* = 0.004). Tukey’s post-hoc comparisons revealed a significant decrease in tail withdrawal latency on P7 and P14 (\**p* < 0.05).

### 3.3. Ultrasonic vocalizations (USVs)

In adult mice, USVs are exclusively emitted during social and mating interactions. However, in mouse pups, USVs are emitted in response to temperature changes and isolation and serve as distress signals to promote maternal attention and care. In examining total USVs on P7, there were no significant main effects (Morphine Treatment: *F*_(1,132)_ = 0.09, *p* = 0.76; Sex: *F*_(1,132)_ = 0.05, *p* = 0.82) and no interaction (*F*_(1,132)_ = 0.89, *p* = 0.35; **Figure 3A**). In examining total USVs on P14, there was again no significant main effects (Morphine Treatment: *F*_(1,108)_ = 2.24, *p* = 0.137; Sex: *F*_(1,108)_ = 0.11, *p* = 0.741) nor was there an interaction (*F*_(1,108)_ = 2.27, *p* = 0.13); **Figure 3B**). In examining the peak frequency on P7, there were no significant main effects (Morphine Treatment: *F*_(1,95)_ = 1.02, *p* = 0.316; Sex: *F*_(1,95)_ = 2.69, *p* = 0.10) and no interaction (*F*_(1,95)_ = 1.36, *p* = 0.25; **Figure 3C**). In examining peak frequency on P14, there was a main effect of Morphine Treatment (*F*_(1,95)_ = 5.37, \**p* = 0.02; MOR > SAL; **Figure 3D**), a near-significant effect of Sex (*F*_(1,95)_ = 3.86, *p* = 0.05), but no interaction (*F*_(1,95)_ = 0.01, *p* = 0.91). Regarding peak amplitude, for P7, there were no significant main effects (Morphine Treatment: *F*_(1,95)_ = 0.61, *p* = 0.44; Sex: *F*_(1,108)_ = 0.61, *p* = 0.44; **Figure 3E**), but a near-significant Morphine Treatment x Sex interaction (*F*_(1,95)_ = 3.89, *p* = 0.05). For peak amplitude on P14, there was a main effect of Morphine Treatment (*F*_(1,95)_ = 21.56, **\*\*\****p* < 0.001; MOR < SAL; **Figure 3F**), but no effect of Sex (*F*_(1,95)_ = 0.74, *p* = 0.39) and no interaction (*F*_(1,95)_ = 0.60, *p* = 0.44). In examining the time dependency of USVs across 1-min time bins on P7, Mauchly’s Test indicated the assumption of sphericity had been violated (χ^2^_44_ = 150.05, \*\*\**p* < 0.001), so the Greenhouse-Geisser correction was applied (ε = 0.76). Repeated measures ANOVA (Time, Morphine Treatment, Sex) indicated a main effect of Time (*F*_(6.8,895.6)_ = 7.16, \*\*\**p* < 0.001), a Morphine Treatment x Time x morphine interaction (*F*_(6.8,895.6)_ = 3.91, \*\*\**p* < 0.001), and a Sex x Time interaction (*F*_(6.8,895.6)_ = 2.06, \**p* = 0.047). Post-hoc multiple comparisons revealed that morphine-treated pups emitted more USVs than saline-treated pups during the second minute of testing (Bonferroni adjusted **\****p* = 0.05 **Figure 3G**). In examining USVs across 1-min time bins on P14, Mauchly’s Test indicated the assumption of sphericity had been violated (χ^2^_104_ = 444.45, \*\*\**p* < 0.001), so the Greenhouse-Geisser correction was applied (ε = 0.57). There was a main effect of Time (*F*_(8.0,855.45)_ = 74.22, \*\*\**p* < 0.001), a Sex x Time interaction (*F*_(8.0,855.45)_ = 2.06, \**p* = 0.04), but no effect of Morphine Treatment (*F*_(1,107)_ = 2.69, *p* = 0.10) nor interactions with Morphine Treatment (all *p* > 0.41; **Figure 3H**).

**Figure 3.**
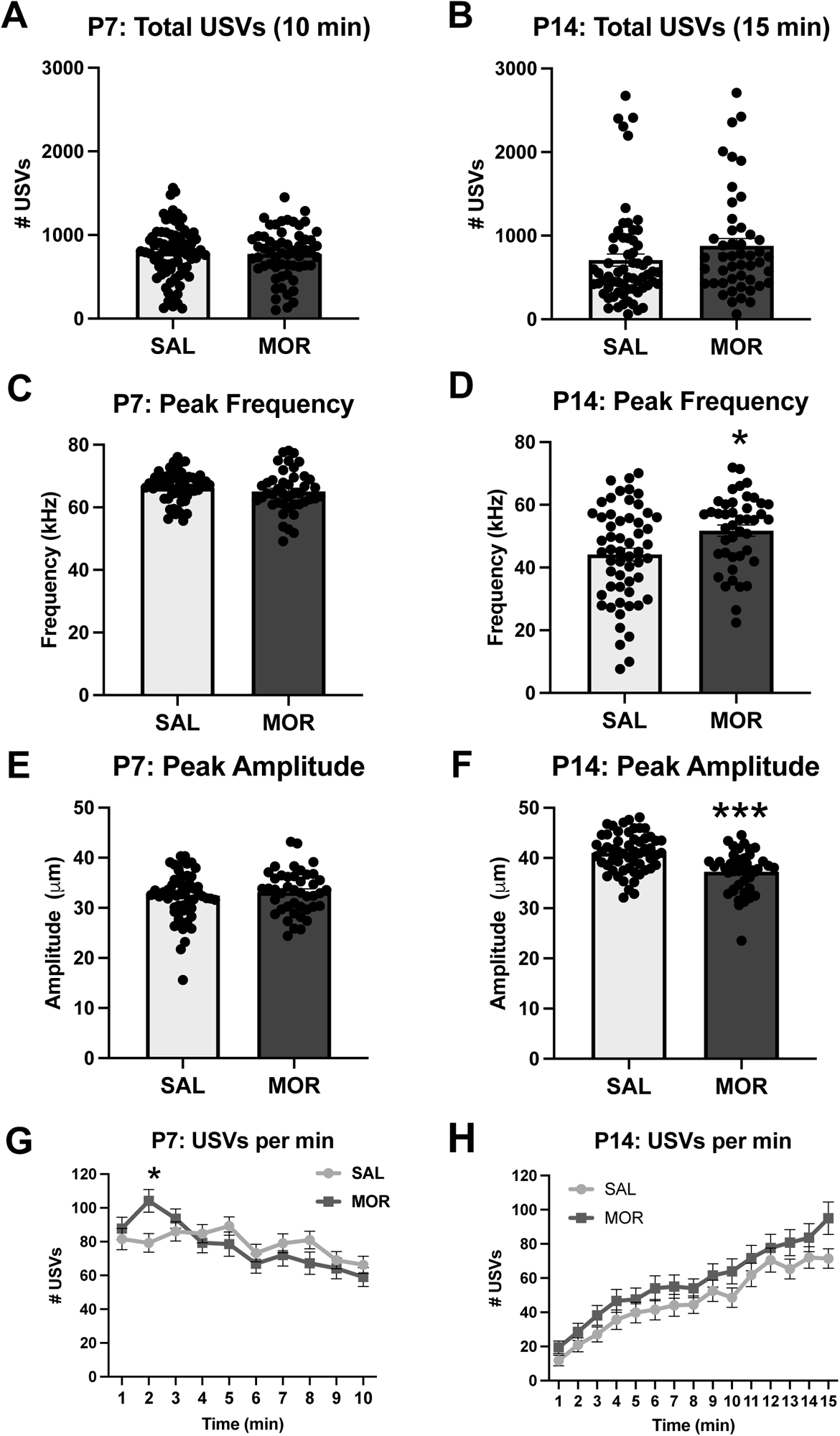
USV emissions and features during spontaneous withdrawal on P7 and P14 following once daily morphine exposure (15 mg/kg, s.c.) from P1-P14. Data are plotted as the mean ± SEM. **A-B.** P7, P14 total USVs: There were no significant main effects of Morphine Treatment and Sex and no interactions. SAL P7: n = 77 (31 females, 46 males); SAL P14: n = 64 (25 females, 39 males); MOR P7: n = 59 (37 females, 22 males); MOR P14: n = 48 (30 females, 18 males). **C.** P7 peak frequency: No significant main effects nor interactions. **D.** P14 peak frequency: Main effect of Morphine Treatment (\**p* = 0.02; MOR > SAL) but no effect of Sex and no interaction. **E.** P7 peak amplitude: No effect of Morphine Treatment, Sex, or interaction. **F.** P14 peak amplitude: Main effect of Morphine Treatment (\*\*\**p* < 0.001; MOR < SAL) but no effect of Sex and nor interaction. SAL P7 – P14: n = 56 (22 females, 34 males); MOR P7 – P14: n = 43 (25 females, 18 males). **G.** Normalized P7 USVs per min: Main effect of Time (\*\*\**p* < 0.001) and a Morphine Treatment x Time interaction (\**p* = 0.047). Morphine treated pups emitted a greater number of USVs than saline treated pups during the second minute of testing (\**p*_adjusted_ = 0.03). **H.** Normalized P14 USVs per min: Main effect of Time (\*\*\**p* < 0.001) and a Sex x Time interaction (\**p* = 0.04). SAL P7: n = 77 (31 females, 46 males); SAL P14: n = 64 (25 females, 39 males); MOR P7: n = 59 (37 females, 22 males); MOR P14: n = 48 (30 females, 18 males).

### 3.4. Locomotor activity during USV recordings

During the withdrawal state, morphine-treated pups displayed increased locomotor activity during USV recordings on P7 compared to saline-treated controls (**Figure 4A**) but not on P14 (**Figure 4B**). Distance traveled on P7 was normalized prior to analysis (resulting Shapiro-Wilk p = 0.99). On P7, there was a main effect of Morphine Treatment (*F*_(1,98)_ = 12.63, \*\*\**p* < 0.001) but no effect of Sex (*F*_(1,98)_ = 0.32, *p* = 0.57) and no significant Morphine Treatment x Sex interaction (*F*_(1,98)_ = 0.412, *p* = 0.52). On P14, there were no significant main effects (Morphine Treatment: *F*_(1,111)_ = 0.11, *p* = 0.74; Sex: *F*_(1,111)_ = 1.09, *p* = 0.30) or interactions (*F*_(1,111)_ = 1.27, *p* = 0.26).

**Figure 4.**
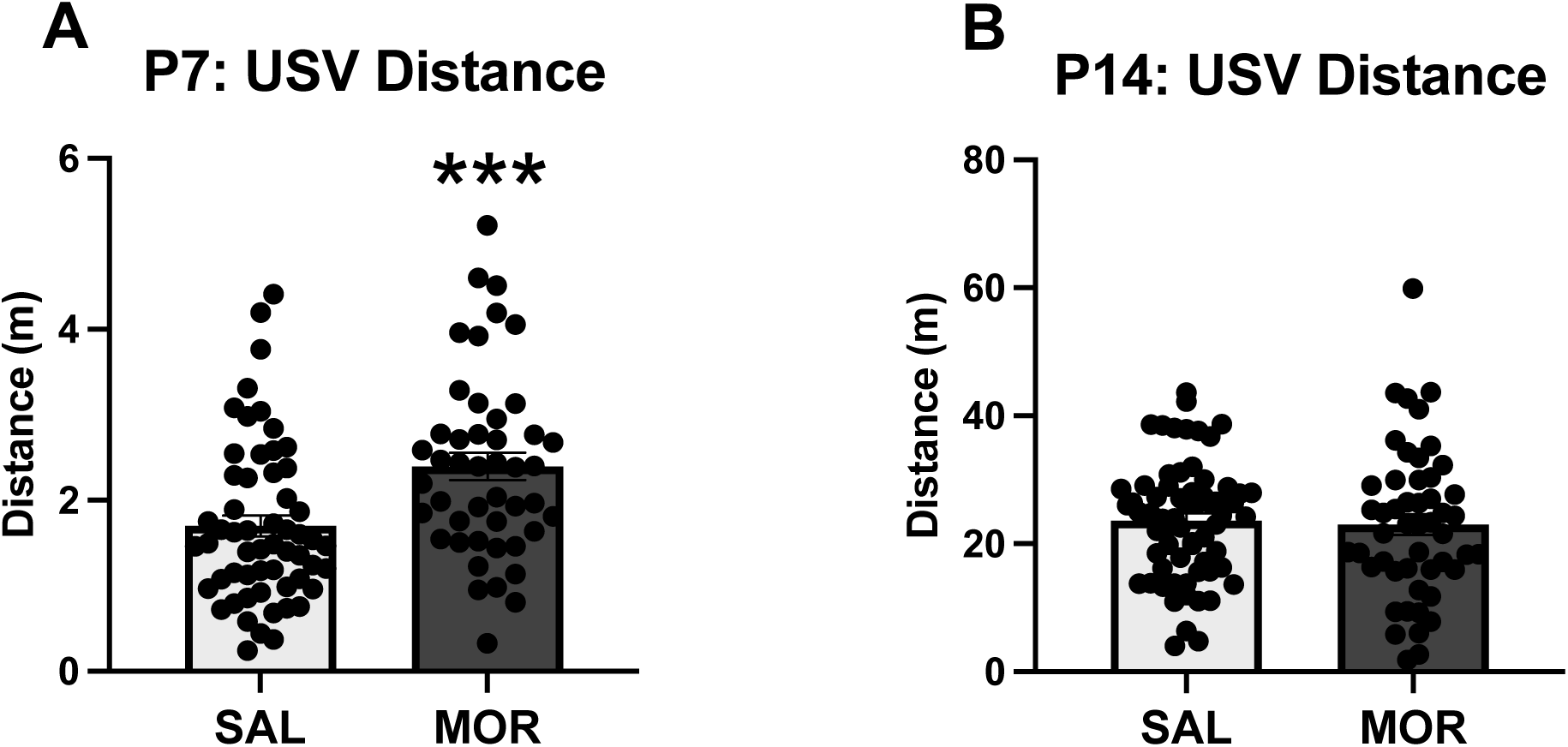
Locomotor activity during spontaneous withdrawal during USV recordings on P7 and P14. Data are plotted as the mean ± SEM. **A.** P7 distance: Main effect of Morphine Treatment (\*\*\**p* < 0.001) but no effect of Sex and no interaction. SAL P7, n = 57 (24 females, 33 males); MOR P7, n = 46 (29 females, 17 males). **B.** P14 distance: No main effects or interactions. SAL P14: n = 65 (24 females, 41 males); MOR P14: n = 50 (30 females, 20 males).

### 3.5. Transcriptome analysis of nucleus accumbens on P15 during spontaneous morphine withdrawal

We identified a total of 17 genes with an adjusted *p* < 0.05 (**Table S1**). Of these genes, 13 showed a log_2_fold-change (FC) ≤ -0.26 (1.2-fold), corresponding to at least a 20% change in gene expression. The genes are labelled in a volcano plot showing gene expression relative to saline-treated controls (**Figure 5A**). Notably, all 13 of these genes were downregulated in morphine-treated pups. A total of 90 downregulated genes with an unadjusted *p* < 0.01 and log_2_FC ≤ -0.26 were used for the enrichment analysis. The top enriched biological processes included myelination (17), ensheathment of neurons and axons (9), oligodendrocyte differentiation (7) and gliogenesis (5) (**Figure 5B, Table S2**). No enriched terms were associated with upregulated differentially expressed genes (log_2_FC > 0.26 and *p* < 0.01). For the complete list of all differentially expressed genes (308; *p* < 0.01), including 105 upregulated genes and 203 downregulated genes see **Table S1**.

**Figure 5.**
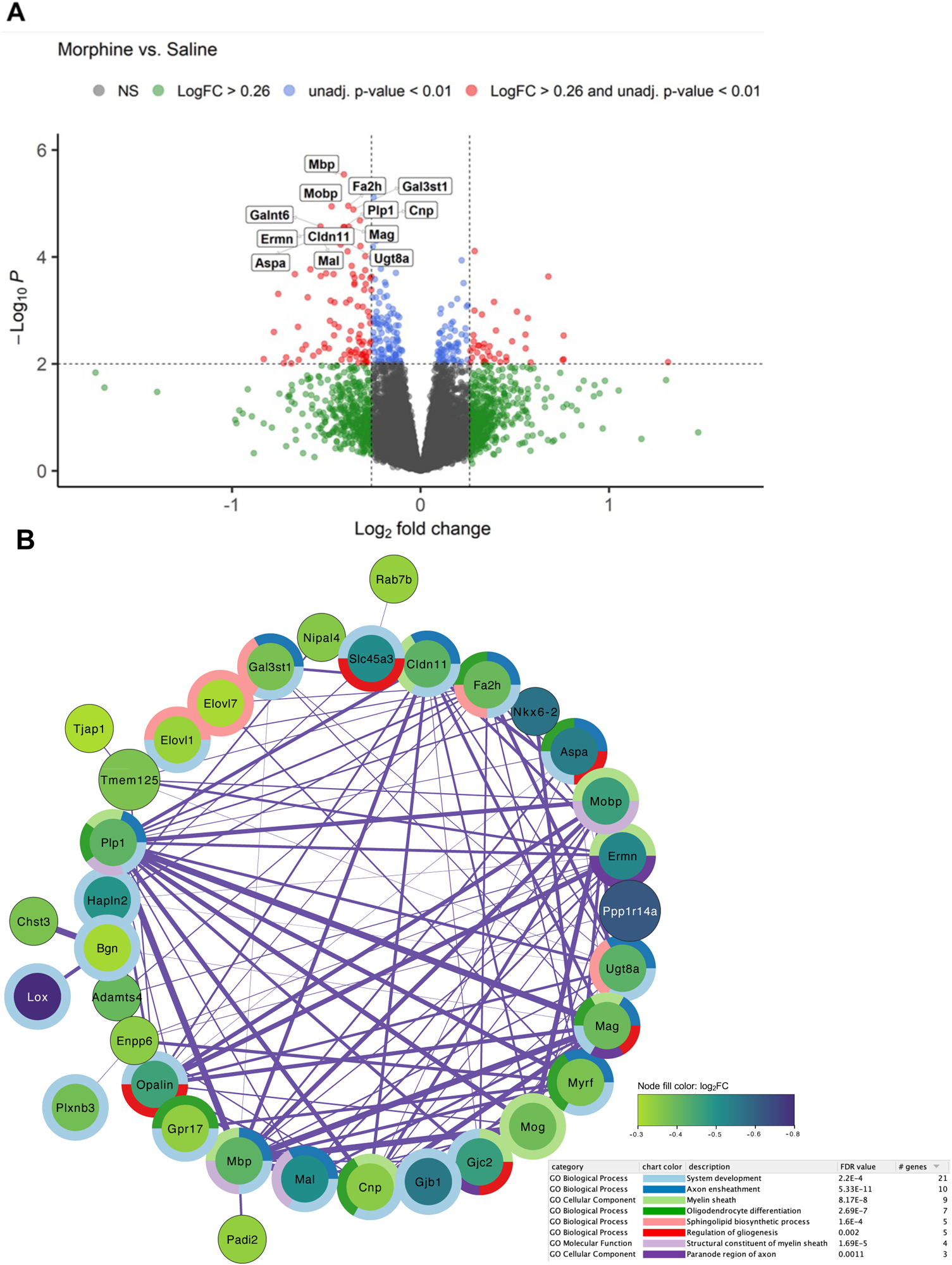
Bulk RNA-seq analysis in the nucleus accumbens (NAc) during spontaneous opioid withdrawal on P15 revealed significant down-regulation of myelin- and oligodendrocyte-related genes. **A.** Volcano plot of gene expression in morphine mice [n = 14; 6 females, 8 males)] relative to saline mice [n = 14 (6 females, 8 males)]. The 13 genes with log_2_FC ≤ -0.26 and adjusted p values < 0.05 are labeled. Genes (n = 90) with log_2_FC ≤ -0.26 and unadjusted p < 0.01 were used as input for gene enrichment and network analysis. **B.** The largest subnetwork plot of significantly downregulated genes following neonatal morphine exposure is shown. Node color depicts gene log_2_FC from -0.3 (purple) to -0.8 (green) relative to saline controls. Surrounding donut plots show the top 8 significantly enriched GO pathways. Connecting lines represent gene interactions, with stronger interactions depicted as thicker lines. See **Supplementary Table 1** for the full DEG list with unadjusted p < 0.01, and **Supplementary Table 2** for a list of significantly enriched GO Biological Process pathways associated with morphine treatment,

### 3.6. Spatial memory (Barnes Maze)

The long-term effects of neonatal opioid exposure on adolescent and adult behavioral outcomes are poorly understood. Here, we compared the spatial memory of young adult morphine-exposed mice to saline mice using the Barnes maze. These measures represent the probe day (48 h after final training sessions). The % time in the target zone and primary latency were normalized prior to analysis (resulting Shapiro-Wilk statistics *p* = 0.96 and *p* = 0.82, respectively). There were no significant main effects nor interactions on any of the probe day measures for the Barnes maze test (**Figure 6**), indicating no long-term effects on spatial memory as assessed in this test following this specific neonatal morphine regimen. For % time in target zone, there were no significant main effects (Morphine Treatment: *F*_(1,52)_ = 2.53, *p* = 0.12; Sex: *F*_(1,52)_ = 0.010, *p* = 0.92) and no interaction (*F*_(1,52)_ = 0.060, *p* = 0.81; **Figure 6A**). For primary latency, there were no significant main effects (Morphine Treatment: *F*_(1,52)_ = 0.112, *p* = 0.74; Sex: *F*_(1,95)_ = 1.21, *p* = 0.28) and no interaction (*F*_(1,52)_ = 0.23, *p* = 0.64; **Figure 6B**). For mean distance to target hole, there were no significant main effects (Morphine Treatment: *F*_(1,52)_ = 1.11, *p* = 0.30; Sex: *F*_(1,52)_ = 2.134, *p* = 0.15) and no interaction (*F*_(1,52)_ = 0.04, *p* = 0.85; **Figure 6C**). For entries to the target hole, there were no significant main effects (Morphine Treatment: *F*_(1,52)_ = 0.092, *p* = 0.76; Sex: *F*_(1,52)_ = 2.44, *p* = 0.12) and no interaction (*F*_(1,52)_ = 0.627, *p* = 0.43; **Figure 6D**).

**Figure 6.**
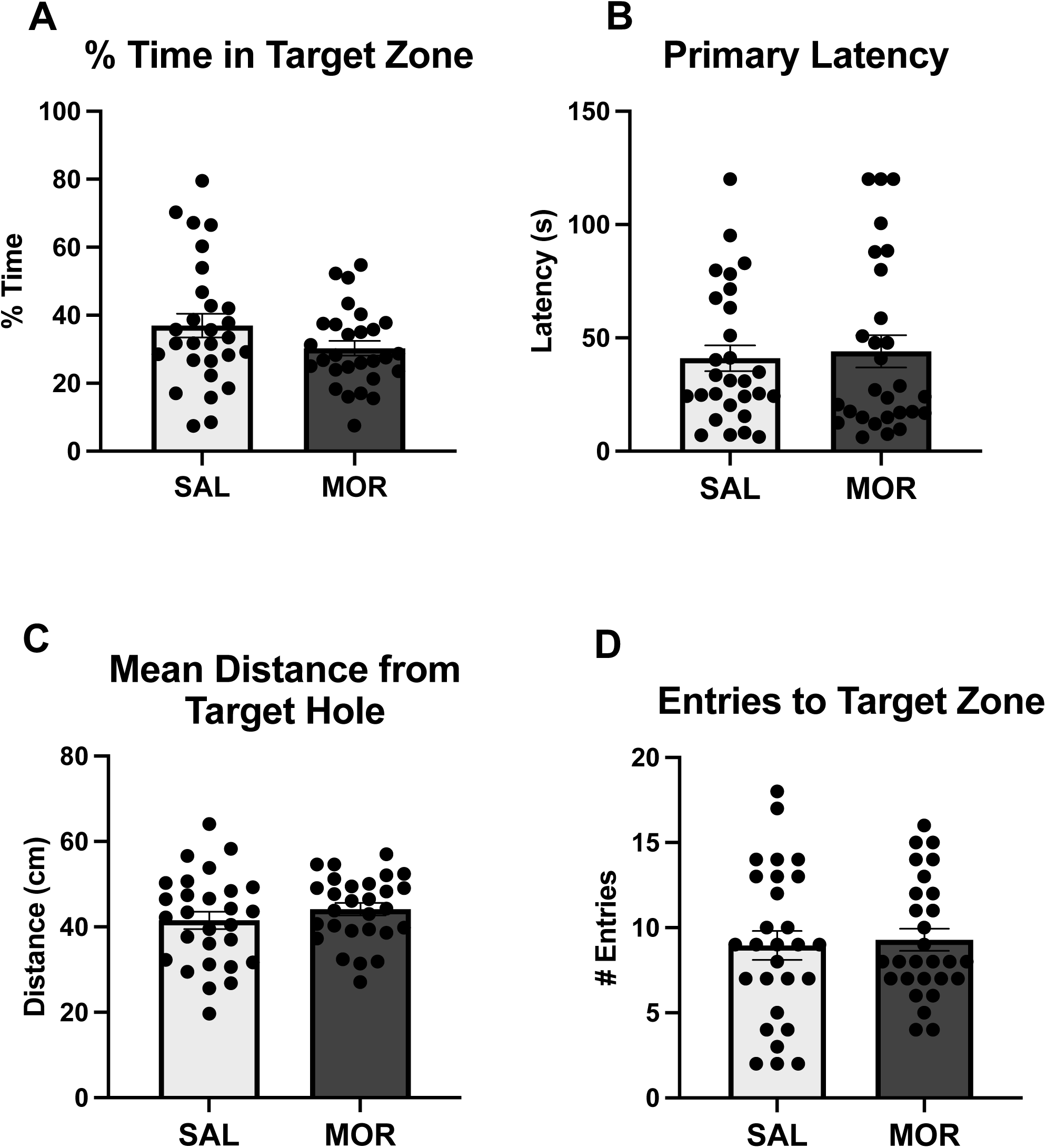
No change in adult spatial learning as measured via the Barnes assay following once daily morphine exposure (15 mg/kg, s.c.) from P1-P14. Data are plotted as the mean ± SEM. **A-D.** % time in the target zone, primary latency to identify the target hole, mean distance to the target hole, and entries to the target zone. No effect of Morphine Treatment, Sex, or interactions SAL: n = 28 (14 females, 14 males); MOR: n = 28 (16 females, 12 males).

### 3.7. Fear conditioning

We also assessed associative learning and memory using Palovian fear conditioning. Both saline- and morphine-exposed mice showed similar fear conditioning freezing behavior, as presented as percent time freezing, on Day 1 – 3 (**Figure 7**), indicating that prior morphine exposure did not impair learning and memory in young adults in this task. Specifically, there was no significant effect of Morphine Treatment on % time freezing on Day 1 (*F*_(1,56)_ = 0.72, *p* = 0.40; **Figure 7A**), Day 2 (*F*_(1,56)_ = 0.17, *p* = 0.69; **Figure 7B**), or Day 3 (*F*_(1,56)_ = 0.00, *p* = 0.99; **Figure 7C**) of fear conditioning tests. There were no significant interactions with Morphine Treatment and sex on Day 1 (*F*_(1,56)_ = 0.002, *p* = 0.96), Day 2 (*F*_(1,56)_ = 0.725, *p* = 0.40), or Day 3 (*F*_(1,56)_ = 0.013, *p* = 0.91).

**Figure 7.**
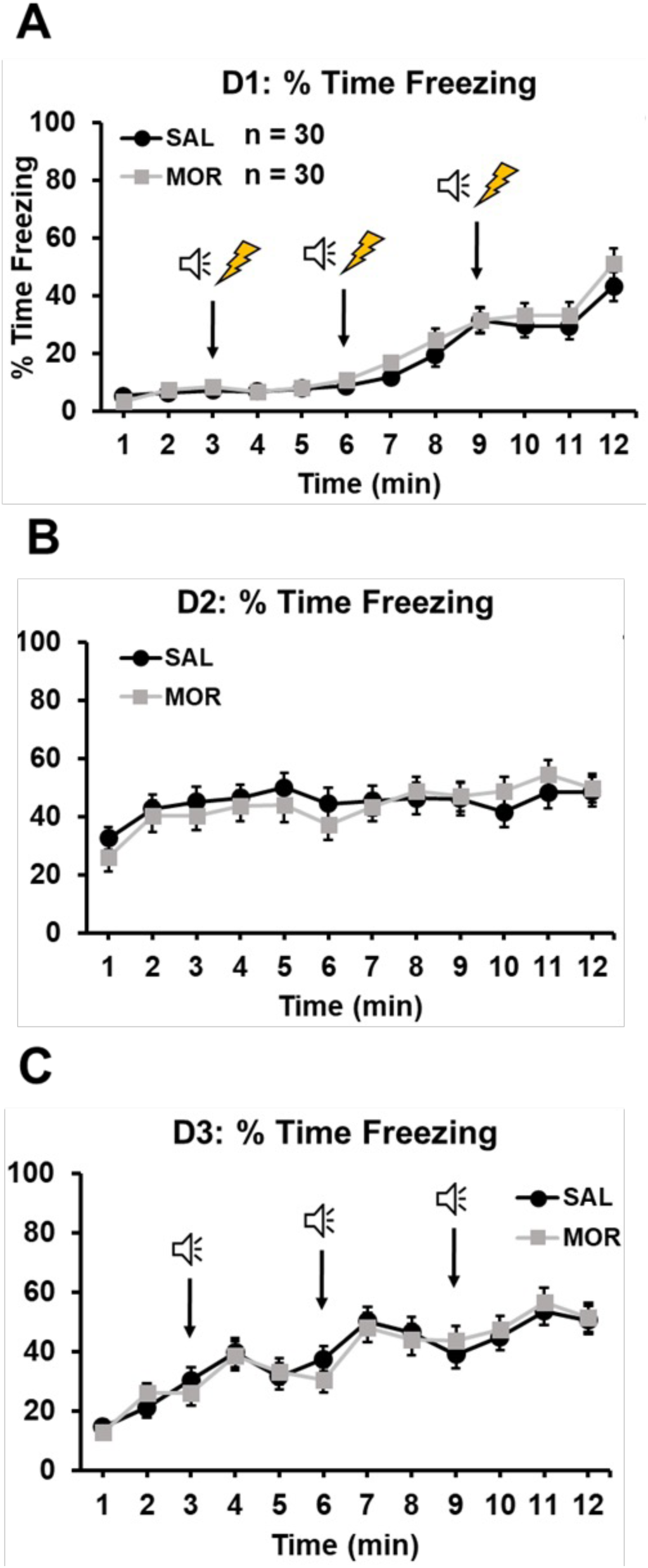
No change in on adult conditioned freezing behavior following daily morphine exposure (15 mg/kg, s.c.) from P1-P14. Arrows in panel A indicate onset of the tone & subsequent foot-shock. In panel C, arrows indicate presentation of the conditioned stimulus (tone). **A-C.** No effect of Morphine Treatment or interactions with Day on % time freezing on Day 1, Day 2, or Day 3 of the fear conditioning tests. Main effect of time: (panel A). Main effect of time (panel B). Main effect of Day (panel C). Data are plotted as the mean ± SEM. SAL, n = 30 (14 females, 16 males); MOR, n = 30 (18 females, 12 males).

### 3.8. ICSS and methamphetamine potentiation of brain stimulation reward

Next, we explored the effects of postnatal morphine exposure on brain stimulation reward via intracranial self-stimulation (**ICSS**) to the medial forebrain bundle containing mesolimbic dopaminergic reward fibers of passage. There was no effect of postnatal Morphine Treatment on ICSS rewards delivered (**Figure 8A-B**) or responses for stimulation (**Figure 8C-D**) following reward potentiation with methamphetamine (MA). Specifically, there was no effect of Morphine Treatment on total rewards delivered (*F*_(1,18)_ = 0.13, *p* = 0.73; **Figure 8A**), total rewards delivered as % vehicle (*F*_(1,18)_ = 2.99, *p* = 0.10; **Figure 8B**), total responses for stimulation (*F*_(1,18)_ = 0.24, *p* = 0.63; **Figure 8C**), or total responses for stimulation as % vehicle responses (**Figure 8D**; *F*_(1,18)_ = 3.21, *p* = 0.09). However, methamphetamine effectively potentiated reward, as there was a significant effect of Methamphetamine Dose on both total rewards (*F*_(1,18)_ = 4.77, *p* = 0.001; **Figure 8A**) and total responses (*F*_(1,18)_ = 3.77, *p* = 0.007; **Figure 8C**). There were no significant MA Dose x Morphine Treatment interactions for any of the measures (all *p* ≥ 0.47). These negative results suggest negligible long-term effects of once-daily, 3^rd^-trimester-equivalent morphine exposure on reward facilitation by substances with high abuse potential.

**Figure 8.**
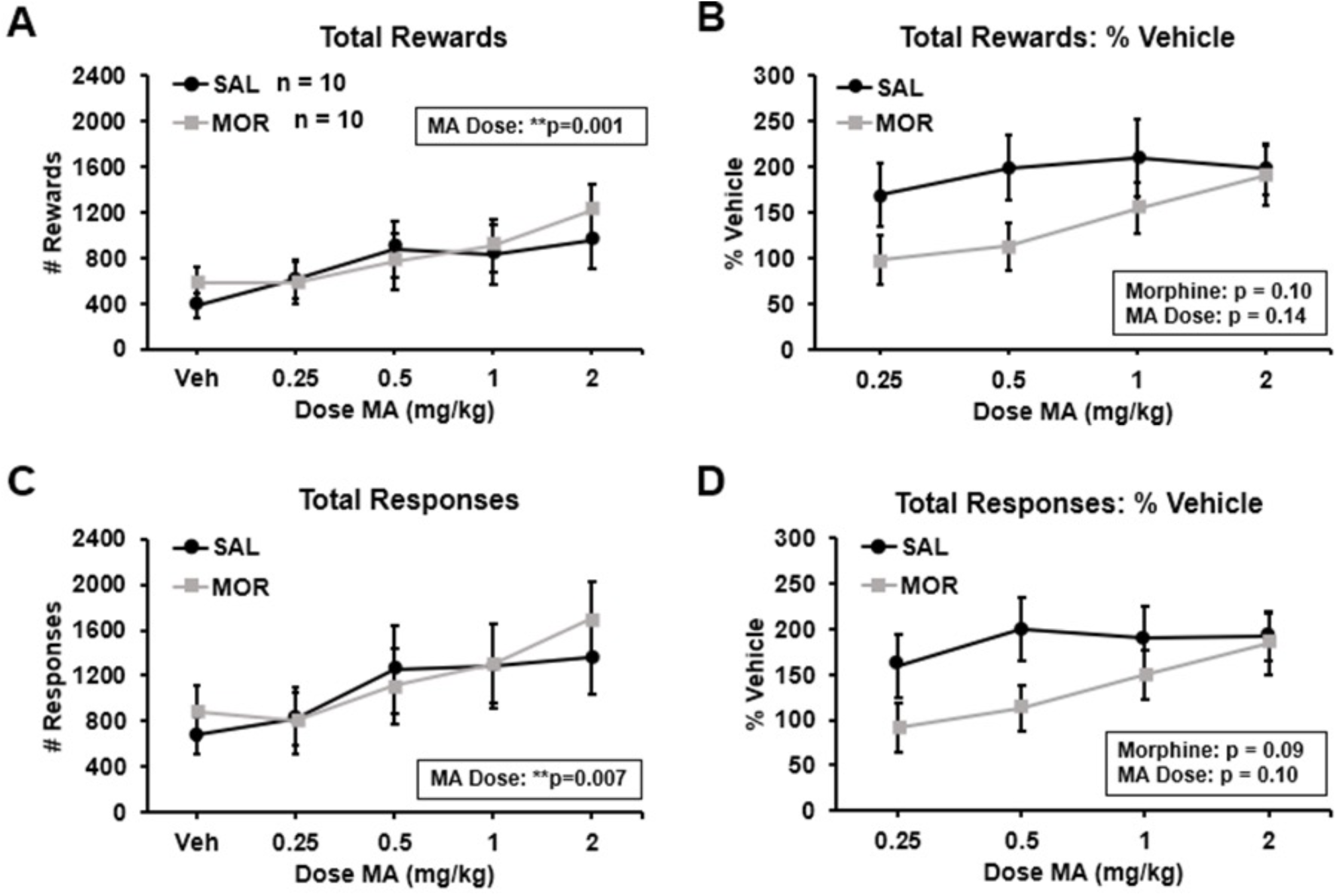
No changes in adult methamphetamine potentiation of brain stimulation reward as measured via intracranial self-stimulation (ICSS) following once daily morphine (15 mg/kg, s.c.) from P1-P14. **A-D.** Total rewards delivered, total rewards delivered as % vehicle, total responses for stimulation, total responses for stimulation as % vehicle, respectively. No effect of Morphine Treatment or interactions with Methamphetamine Dose for any of these measures. There was a significant effect of Methamphetamine Dose on total rewards (**A**: \*\*\**p* = 0.001) and total responses (**C**: \*\**p* = 0.007), indicating methamphetamine-induced potentiation of brain stimulation reward. Data are plotted as the mean ± SEM. Sample sizes are shown in panel A next to the figure legend. Sample size by Sex: SAL: n = 10 (6 females, 4 males); MOR: n = 10 (7 females, 3 males).

## 4. Discussion

We used a third trimester-approximate model of once daily morphine exposure to induce NOWS model traits in outbred CFW mice, with the goal of increasing pup survival compared to our prior twice daily model (100% vs ∼85%; Borrelli et al., 2021) and then examining the effect of P1-P14 exposure in adult memory and reward function. We replicated some of the NOWS model traits that we previously reported, including robust thermal hyperalgesia on P7 and P14, time-dependent differences in USV emissions on P7, alterations to USV features (decreased mean amplitude and increased mean frequency) on P14, and increased locomotor activity on P7 during USV testing on P7. These findings are similar to our previous results where we observed withdrawal-like phenotypes in a model of *twice*-daily morphine administration (15.0 mg/kg twice daily from P1-14) (Borrelli, Yao, et al. 2021). While our previous study examined brainstem transcriptomic adaptations, the current study explored transcriptomic adaptations within the NAc following the third trimester model of morphine exposure. We identified 17 genes that met the genome-wide threshold for statistical significance; all of these genes were downregulated in morphine-exposed pups. Exploratory enrichment analysis of the larger downregulated gene set (*p* < 0.01, unadjusted; absolute fold-change of 1.2-fold or greater) identified pathways that included neuron and axon ensheathment, myelination and oligodendrocyte differentiation. Several of these genes comprising these enriched pathways surpassed the adjusted significance threshold for differential gene expression (*p*_adjusted_ < 0.05), including *Mbp*, *Mal*, *Plp1*, and *Gal3st1* (“myelination”), and *Aspa*, *Plp1*, *Fa2h*, *Cnp*, and *Mag* (“oligodendrocyte differentiation”).

How might morphine affect oligodendrocyte gene expression? Both μ and κ opioid receptors are expressed on oligodendrocytes and have been shown to play a role in the maturation and differentiation of these cells (Eschenroeder et al. 2012; Mohamed et al. 2020; Velasco, Mohamed, and Sato-Bigbee 2021). Thus, activation of the mu opioid receptor directly on oligodendrocytes could modulate their function through Gi/Go signaling, disrupt synaptic connectivity during brain development. Alternatively, morphine could affect myelination/oligodendrocyte function indirectly by binding to opioid receptors on neurons or neural progenitor cells. Several studies reported disruption of myelin-related gene expression in the brain following perinatal exposure to opioids, and these effects vary depending on the opioid drug and dose (Vestal-Laborde et al. 2014; Jantzie et al. 2020; Oberoi et al. 2019). For example, upregulation of myelin-related proteins (MBP PLP, MOG) has been observed in mouse pups at P11 and P19 in whole-brain homogenate following perinatal methadone exposure (9 mg/kg/day maternal minipump administration from gestational day 7 – P21; postnatal exposure via maternal milk) (Vestal-Laborde et al. 2014). In contrast, perinatal methadone (12 mg/kg/day maternal minipump administration from gestational day 16 – P21; postnatal exposure via maternal milk) has also been shown to result in decreased expression of MBP in the cerebral cortex of mouse pups at P21 (Jantzie et al. 2020). Another study provided drinking water containing 150 mg/ml methadone to pregnant rat dams from gestational day 7 through either P7 or P19. This study found down-regulation of MBP and PLP at both P7 and P19 in the brainstem, cerebral cortex, and hippocampus of methadone-exposed rat pups (Oberoi et al. 2019). Additionally, some studies have shown morphine-induced alterations in related genes, such as *Fa2h* and *Plp1*, which are associated with neurodegeneration (Hardt et al., 2020; Neff et al., 2021; Potter et al., 2011).

We found that once daily perinatal morphine administration (15 mg/kg, s.c.) in the third trimester-approximate model did not result in significant long-term changes in adult behavioral measures of cognition, learning and memory, and reward function, including the Barnes maze assessment of spatial memory, emotional learning via fear conditioning, and baseline and methamphetamine-facilitated ICSS thresholds. Perinatal opioid exposure has previously been reported to lead to long-term disruptions in spatial memory function; including decreased performances in the Barnes maze, Morris water maze, y-maze and 8-arm radial maze tasks (Martin et al. 2021; Wang and Han 2009; Niu et al. 2009; Lin et al. 2009). There is also evidence that prenatal exposure to morphine (from gestational day 9-18) impairs acquisition and delays extinction of contextual fear learning during adulthood in rats (Tan et al. 2015). Tan *et al*. propose that prenatal morphine exposure inhibits both long-term potentiation (LTP) and long-term depression (LTD) in the CA1 subregion of the hippocampus, inhibiting both acquisition and extinction of contextual fear learning (Tan et al. 2015). Other work indicates that prenatal morphine exposure can impair both the induction and maintenance of LTP in the hippocampus of adult rats (Villarreal, Derrick, and Vathy 2008; Velísek et al. 2000). These studies all implemented prenatal opioid exposure (heroin, oxycodone, or morphine) to either rat or mouse pups that overlapped with the gestational developmental correlate of the first and second human trimesters, suggesting that opioid exposure during the first two trimesters may be more critical for inducing long-term neurodevelopmental effects on adult brain function. It should also be noted that the regimen in the current study was quite mild, with only once daily injections of a moderate dose of morphine (15 mg/kg, s.c.) and was associated with a less severe withdrawal behavioral repertoire compared to twice daily injections (Borrelli, Yao, et al., 2021). Thus, it is possible that with more frequent dosing during the third trimester-approximate period, significant long-term neurobehavioral effects might be observed in adulthood.

We did not identify significant alterations in reward facilitation following MA treatment during adulthood in mice perinatally exposed to morphine. Gestational opioid exposure has previously been linked to alterations in MA locomotor sensitization (Chiang, Hung, and Ho 2014; Wong et al. 2014), MA intravenous self-administration (Shen et al. 2016), and MA-paired conditioned place preference (Chiang, Hung, and Ho 2014). The lack of significant alterations in MA-facilitated ICSS behaviors could be attributed to our small sample size. It should also be noted that the regimen in the current study was quite mild, with only once daily injections of a moderate dose of morphine (15 mg/kg, s.c.) and was associated with a less severe withdrawal behavioral repertoire compared to twice daily injections (Borrelli, Yao et al., 2021). Thus, it is possible that with more frequent dosing during the third trimester-approximate period, significant long-term neurobehavioral effects might be observed in adulthood. Future NOWS studies will involve more robust treatment regimens for third trimester-approximate exposure (twice daily with 10 mg/kg s.c. morphine) and larger sample sizes for neurobehavioral measurements in adulthood.

## Conclusions

To summarize, our once daily morphine regimen during the approximate third trimester period induced less robust signs of NOWS model traits compared to our twice daily regimen and did not significantly affect long-term neurobehavioral outcomes on learning or reward sensitivity in adult CFW mice. Nevertheless, we identified a NAc transcriptional profile predictive of reduced myelination and oligodendrocyte differentiation following this exposure that warrants further functional testing. This transcriptional profile was quite different from what we identified in the brainstem using the twice daily protocol (Borrelli, Yao, et al., 2021), suggesting that the effects on the myelination may be brain region-specific or that white matter dysfunction is more readily detectable in the NAc. Given the accumulating evidence for myelin disruption following perinatal opioid exposure, it will be important to determine the persistence and functional relevance of these changes during adolescence and adulthood and how they relate to neurobehavioral function.

## CRediT authorship contribution

**Kristyn N. Borrelli:** Conceptualization, Data curation, Methodology, Formal analysis, Funding acquisition, Investigation, Validation, Writing – original draft, Writing – review & editing, Visualization. **Kelly K. Wingfield:** Formal analysis, Writing – review & editing, Visualization. **Emily J. Yao:** Data curation, Methodology, Formal analysis, Investigation, Validation. **Catalina A. Zamorano:** Investigation, Formal analysis, Validation, Visualization. **Katherine N. Sena:** Investigation, Formal analysis, Validation, Visualization. **Jacob A. Beierle:** Investigation, Data curation, Formal analysis, Validation, Visualization. **Michelle A. Roos:** Investigation, Formal analysis, Validation, Visualization. **Huiping Zhang:** Writing – review & editing. **Elisha M. Wachman:** Writing – review & editing. **Camron D. Bryant:** Conceptualization, Writing – original draft, Writing – review & editing, Visualization, Supervision, Funding acquisition

## Data availability

Differential gene expression data is available on the Gene Expression Omnibus (https://www.ncbi.nlm.nih.gov/geo/query/acc.cgi?acc=GSE239919). Additional data will be made available upon request.

## Acknowledgements

We would like to thank the Genome Science Institute at Boston University, NIH/NIDA U01DA050243, T32GM008541, Boston University’s Undergraduate Research Opportunities Program (UROP), and the animal care staff at the Boston University Animal Science Center.

