## Supplemental Tables for "Decreased myelin-related gene expression in the nucleus accumbens during spontaneous neonatal opioid withdrawal in the absence of long-term behavioral effects in adult outbred CFW mice"

**SUPPLEMENTARY INFORMATION**

**Supplementary Table S1**. Differentially expressed genes (p<0.01) between morphine vs. saline mice.

| **Gene** | **logFC**  **(MOR-SAL)** | **AveExpr**  **(cpm)** | **t** | **P.Value** | **adj.P.Val** |
| --- | --- | --- | --- | --- | --- |
| Mbp | -0.41 | 10.78 | 5.861 | 2.89E-06 | 0.044 |
| Bcas1 | -0.25 | 6.61 | 5.499 | 7.62E-06 | 0.044 |
| Fa2h | -0.38 | 4.71 | 5.357 | 1.12E-05 | 0.044 |
| Mobp | -0.47 | 8.42 | 5.352 | 1.13E-05 | 0.044 |
| Gal3st1 | -0.36 | 3.73 | 5.300 | 1.30E-05 | 0.044 |
| Cnp | -0.32 | 8.47 | 5.127 | 2.08E-05 | 0.047 |
| Galnt6 | -0.53 | 1.41 | 5.029 | 2.71E-05 | 0.047 |
| Mag | -0.38 | 7.67 | 5.026 | 2.73E-05 | 0.047 |
| Plp1 | -0.40 | 10.34 | 5.022 | 2.77E-05 | 0.047 |
| Cldn11 | -0.41 | 6.90 | 5.019 | 2.79E-05 | 0.047 |
| Ermn | -0.55 | 3.75 | 4.962 | 3.25E-05 | 0.049 |
| Tmem229a | -0.20 | 5.58 | 4.916 | 3.68E-05 | 0.049 |
| Sox10 | -0.25 | 5.39 | 4.876 | 4.10E-05 | 0.049 |
| Mal | -0.53 | 6.07 | 4.857 | 4.32E-05 | 0.049 |
| Aspa | -0.57 | 2.03 | 4.819 | 4.78E-05 | 0.049 |
| Ugt8a | -0.41 | 6.65 | 4.812 | 4.88E-05 | 0.049 |
| Gltp | -0.24 | 5.30 | 4.804 | 4.98E-05 | 0.049 |
| Trf | -0.42 | 5.32 | 4.742 | 5.89E-05 | 0.054 |
| Enpp6 | -0.32 | 4.86 | 4.718 | 6.29E-05 | 0.054 |
| Tspan2 | -0.25 | 7.33 | 4.712 | 6.40E-05 | 0.054 |
| 8430429K09Rik | 0.29 | 2.89 | -4.637 | 7.83E-05 | 0.060 |
| Adamts4 | -0.39 | 5.02 | 4.635 | 7.88E-05 | 0.060 |
| Elovl1 | -0.29 | 3.92 | 4.558 | 9.69E-05 | 0.071 |
| Ttyh2 | -0.26 | 4.51 | 4.535 | 0.000103 | 0.072 |
| Parvb | -0.16 | 5.90 | 4.503 | 0.000113 | 0.075 |
| Nsun3 | 0.22 | 3.80 | -4.492 | 0.000116 | 0.075 |
| Sirt2 | -0.19 | 7.82 | 4.468 | 0.000124 | 0.077 |
| Sema4d | -0.23 | 6.61 | 4.452 | 0.000129 | 0.077 |
| Plxnb3 | -0.37 | 4.96 | 4.403 | 0.000147 | 0.085 |
| Kank1 | -0.21 | 5.08 | 4.355 | 0.000168 | 0.093 |
| Gjb1 | -0.58 | 1.41 | 4.347 | 0.000171 | 0.093 |
| Gsn | -0.29 | 6.37 | 4.332 | 0.000178 | 0.093 |
| Hadha | -0.13 | 6.02 | 4.290 | 0.0002 | 0.096 |
| Gm37111 | -0.50 | 1.72 | 4.281 | 0.000204 | 0.096 |
| Opalin | -0.46 | 2.81 | 4.272 | 0.000209 | 0.096 |
| Tmem88b | -0.36 | 4.67 | 4.270 | 0.000211 | 0.096 |
| Ppp1r14a | -0.67 | 1.18 | 4.269 | 0.000211 | 0.096 |
| Slc45a3 | -0.53 | 1.09 | 4.239 | 0.000229 | 0.097 |
| Cldn23 | 0.68 | -0.51 | -4.232 | 0.000233 | 0.097 |
| Gpr17 | -0.31 | 7.71 | 4.230 | 0.000235 | 0.097 |
| Tjap1 | -0.26 | 4.56 | 4.226 | 0.000237 | 0.097 |
| Chst3 | -0.35 | 2.95 | 4.214 | 0.000245 | 0.098 |
| Elovl7 | -0.27 | 4.18 | 4.205 | 0.00025 | 0.098 |
| Gjc3 | -0.24 | 5.24 | 4.190 | 0.000261 | 0.099 |
| Gdf11 | -0.19 | 4.46 | 4.137 | 0.000301 | 0.111 |
| Snx7 | 0.23 | 4.58 | -4.128 | 0.000308 | 0.111 |
| Tmem125 | -0.35 | 2.33 | 4.125 | 0.000311 | 0.111 |
| 2310058D17Rik | -0.29 | 1.80 | 4.110 | 0.000324 | 0.111 |
| Myrf | -0.35 | 5.32 | 4.102 | 0.00033 | 0.111 |
| Reep3 | -0.18 | 5.84 | 4.101 | 0.000331 | 0.111 |
| Lpar1 | -0.30 | 4.46 | 4.018 | 0.000413 | 0.134 |
| Mcam | -0.26 | 4.51 | 4.016 | 0.000415 | 0.134 |
| Inppl1 | -0.22 | 5.27 | 3.986 | 0.00045 | 0.142 |
| Gm44256 | -0.76 | -0.40 | 3.951 | 0.000494 | 0.154 |
| Cers2 | -0.18 | 5.77 | 3.919 | 0.000538 | 0.164 |
| Tgfbr2 | -0.19 | 4.11 | 3.898 | 0.000568 | 0.164 |
| Nkx6-2 | -0.60 | 1.30 | 3.895 | 0.000573 | 0.164 |
| Grin2c | -0.24 | 5.57 | 3.894 | 0.000574 | 0.164 |
| Gpr37 | -0.25 | 4.61 | 3.893 | 0.000576 | 0.164 |
| Riok2 | 0.19 | 4.13 | -3.870 | 0.000611 | 0.171 |
| Mog | -0.37 | 5.56 | 3.858 | 0.000632 | 0.174 |
| Gjc2 | -0.48 | 3.20 | 3.839 | 0.000664 | 0.180 |
| 9930014A18Rik | 0.39 | 2.60 | -3.821 | 0.000697 | 0.183 |
| Gm49322 | -0.46 | 0.47 | 3.812 | 0.000712 | 0.183 |
| Gm8399 | 3.13 | 1.08 | -3.811 | 0.000715 | 0.183 |
| Plekhh1 | -0.39 | 3.75 | 3.809 | 0.000718 | 0.183 |
| Prkcq | -0.22 | 4.15 | 3.802 | 0.000732 | 0.183 |
| Brcc3 | 0.16 | 5.15 | -3.775 | 0.000787 | 0.193 |
| Alkbh3 | 0.25 | 3.26 | -3.771 | 0.000795 | 0.193 |
| Mecom | -0.33 | 1.89 | 3.751 | 0.000838 | 0.196 |
| Plod1 | -0.17 | 4.88 | 3.747 | 0.000847 | 0.196 |
| Pfn4 | 0.25 | 3.29 | -3.746 | 0.000848 | 0.196 |
| Padi2 | -0.31 | 3.33 | 3.743 | 0.000854 | 0.196 |
| Nop58 | 0.11 | 5.78 | -3.711 | 0.00093 | 0.211 |
| Gna12 | -0.15 | 5.94 | 3.706 | 0.000942 | 0.211 |
| Mkrn3 | 0.29 | 4.21 | -3.677 | 0.001018 | 0.224 |
| Dnajb3 | 0.51 | 0.65 | -3.664 | 0.001051 | 0.224 |
| Snx33 | -0.28 | 2.66 | 3.664 | 0.001052 | 0.224 |
| Neu4 | -0.23 | 4.45 | 3.663 | 0.001053 | 0.224 |
| Phlpp1 | -0.14 | 6.39 | 3.632 | 0.001143 | 0.237 |
| Oga | 0.11 | 7.83 | -3.632 | 0.001144 | 0.237 |
| Ddr1 | -0.16 | 6.22 | 3.627 | 0.001158 | 0.237 |
| Ifi30 | 0.33 | 2.08 | -3.616 | 0.001193 | 0.241 |
| Tmcc2 | -0.12 | 7.31 | 3.599 | 0.001246 | 0.249 |
| Adgra2 | -0.24 | 4.19 | 3.580 | 0.001308 | 0.255 |
| Gpr165 | -0.27 | 3.67 | 3.578 | 0.001315 | 0.255 |
| Snap23 | -0.18 | 4.03 | 3.573 | 0.001332 | 0.255 |
| Rab22a | -0.11 | 5.97 | 3.572 | 0.001337 | 0.255 |
| Cnksr3 | -0.21 | 3.84 | 3.564 | 0.001365 | 0.257 |
| Lgi3 | -0.23 | 5.97 | 3.556 | 0.001394 | 0.258 |
| Fam169b | 0.57 | 0.10 | -3.555 | 0.001397 | 0.258 |
| Zbtb7b | -0.25 | 3.09 | 3.545 | 0.001435 | 0.262 |
| Myo6 | -0.17 | 6.69 | 3.519 | 0.001533 | 0.277 |
| Sec14l5 | -0.48 | 2.60 | 3.510 | 0.00157 | 0.280 |
| Notch3 | -0.25 | 5.11 | 3.501 | 0.001607 | 0.284 |
| Alg6 | 0.17 | 3.83 | -3.496 | 0.001629 | 0.285 |
| Fn3k | -0.20 | 4.06 | 3.480 | 0.001694 | 0.288 |
| Per2 | -0.27 | 5.29 | 3.470 | 0.001739 | 0.288 |
| Phldb1 | -0.19 | 6.55 | 3.470 | 0.001741 | 0.288 |
| Prr18 | -0.24 | 4.18 | 3.466 | 0.001759 | 0.288 |
| Pcgf2 | 0.16 | 5.24 | -3.465 | 0.001764 | 0.288 |
| Cdh5 | -0.23 | 3.96 | 3.464 | 0.001766 | 0.288 |
| Tns3 | -0.23 | 6.71 | 3.463 | 0.00177 | 0.288 |
| 4833438C02Rik | -0.46 | -0.37 | 3.453 | 0.001816 | 0.293 |
| Matn4 | -0.21 | 4.16 | 3.448 | 0.00184 | 0.294 |
| Flt1 | -0.18 | 5.69 | 3.438 | 0.001888 | 0.298 |
| Amt | -0.13 | 4.44 | 3.432 | 0.001917 | 0.298 |
| Elovl5 | -0.12 | 6.68 | 3.431 | 0.001922 | 0.298 |
| Foxf2 | -0.32 | 2.24 | 3.429 | 0.001936 | 0.298 |
| Skil | 0.13 | 7.28 | -3.418 | 0.001989 | 0.303 |
| Nde1 | -0.16 | 4.28 | 3.412 | 0.002019 | 0.304 |
| Gm5112 | -0.65 | -0.17 | 3.410 | 0.002029 | 0.304 |
| 9930012K11Rik | -0.44 | 0.78 | 3.404 | 0.002064 | 0.306 |
| Cd46 | 0.24 | 3.83 | -3.385 | 0.002165 | 0.315 |
| Aldh1b1 | 0.16 | 4.55 | -3.382 | 0.00218 | 0.315 |
| Tspan15 | -0.16 | 4.29 | 3.380 | 0.002192 | 0.315 |
| Itga1 | -0.22 | 4.42 | 3.379 | 0.002198 | 0.315 |
| Rasgrp3 | -0.17 | 3.91 | 3.351 | 0.00236 | 0.335 |
| 1110018N20Rik | 0.40 | 1.60 | -3.348 | 0.002377 | 0.335 |
| Armcx5 | 0.18 | 4.74 | -3.343 | 0.002411 | 0.337 |
| Prpf40a | 0.11 | 6.16 | -3.338 | 0.002439 | 0.337 |
| Zmym1 | -0.37 | 3.34 | 3.336 | 0.00245 | 0.337 |
| Atp1a2 | -0.14 | 9.54 | 3.325 | 0.002525 | 0.344 |
| Lox | -0.78 | 0.28 | 3.322 | 0.00254 | 0.344 |
| Fam114a1 | -0.25 | 2.36 | 3.315 | 0.002586 | 0.347 |
| Kank4 | -0.21 | 2.05 | 3.305 | 0.002652 | 0.347 |
| Gpr37l1 | -0.13 | 6.80 | 3.305 | 0.002656 | 0.347 |
| Abi2 | 0.10 | 8.32 | -3.304 | 0.002662 | 0.347 |
| Armc8 | 0.10 | 6.51 | -3.303 | 0.002668 | 0.347 |
| Prr5l | -0.32 | 2.07 | 3.287 | 0.002778 | 0.358 |
| Rbm12 | 0.22 | 4.39 | -3.280 | 0.002826 | 0.362 |
| Gm14549 | 0.56 | -0.98 | -3.276 | 0.002858 | 0.363 |
| Gm26660 | 0.28 | 2.05 | -3.269 | 0.002908 | 0.364 |
| Klhdc7a | -0.46 | 2.34 | 3.266 | 0.002928 | 0.364 |
| Gm49602 | 0.24 | 3.17 | -3.266 | 0.002931 | 0.364 |
| Col8a2 | 0.76 | -0.07 | -3.263 | 0.002955 | 0.364 |
| Rimklb | 0.16 | 6.28 | -3.247 | 0.003074 | 0.376 |
| Nid1 | -0.25 | 5.08 | 3.239 | 0.00314 | 0.382 |
| Llgl1 | -0.13 | 6.18 | 3.228 | 0.003222 | 0.389 |
| Nipal4 | -0.33 | 2.66 | 3.220 | 0.003294 | 0.394 |
| Prmt9 | 0.12 | 5.01 | -3.217 | 0.003318 | 0.394 |
| Kank2 | -0.22 | 3.27 | 3.214 | 0.003337 | 0.394 |
| Kif1c | -0.12 | 6.05 | 3.197 | 0.003484 | 0.403 |
| Serinc5 | -0.15 | 6.81 | 3.195 | 0.003509 | 0.403 |
| Erbb3 | -0.38 | 3.43 | 3.194 | 0.003517 | 0.403 |
| Myorg | -0.18 | 5.26 | 3.194 | 0.003518 | 0.403 |
| Arhgap29 | -0.24 | 4.65 | 3.192 | 0.003531 | 0.403 |
| Hexdc | 0.24 | 4.87 | -3.178 | 0.003654 | 0.414 |
| Mxd4 | -0.12 | 6.26 | 3.170 | 0.003729 | 0.417 |
| Slc7a2 | -0.21 | 5.17 | 3.168 | 0.003756 | 0.417 |
| Gm42600 | 0.49 | -0.38 | -3.164 | 0.003788 | 0.417 |
| Vrk1 | 0.28 | 4.63 | -3.161 | 0.003817 | 0.417 |
| Sod3 | -0.26 | 2.56 | 3.160 | 0.003829 | 0.417 |
| Slc44a1 | -0.13 | 6.26 | 3.158 | 0.003851 | 0.417 |
| Ldlrap1 | -0.30 | 2.41 | 3.157 | 0.003855 | 0.417 |
| Plekhg3 | -0.32 | 3.46 | 3.153 | 0.003893 | 0.419 |
| Ninl | 0.20 | 3.21 | -3.142 | 0.004001 | 0.426 |
| Zfr | 0.10 | 8.64 | -3.141 | 0.004012 | 0.426 |
| Rapgef3 | -0.27 | 4.16 | 3.130 | 0.004125 | 0.434 |
| Hnmt | 0.19 | 4.63 | -3.124 | 0.004185 | 0.434 |
| Slc13a3 | -0.24 | 4.17 | 3.124 | 0.004192 | 0.434 |
| Clic4 | -0.17 | 6.48 | 3.121 | 0.00422 | 0.434 |
| Hapln2 | -0.51 | 0.54 | 3.119 | 0.004241 | 0.434 |
| Gm11613 | 0.34 | 2.81 | -3.119 | 0.004245 | 0.434 |
| Tsn | 0.10 | 6.67 | -3.114 | 0.004291 | 0.436 |
| Prim1 | 0.20 | 2.78 | -3.111 | 0.004325 | 0.437 |
| Bfsp2 | -0.31 | 1.86 | 3.107 | 0.004372 | 0.437 |
| Dcaf13 | 0.15 | 5.34 | -3.106 | 0.004378 | 0.437 |
| Cep104 | -0.15 | 5.47 | 3.101 | 0.004429 | 0.437 |
| H2bc6 | -0.61 | 0.12 | 3.101 | 0.004432 | 0.437 |
| Zranb1 | 0.15 | 6.33 | -3.092 | 0.004536 | 0.442 |
| Kdm4d | 0.31 | 0.90 | -3.092 | 0.004536 | 0.442 |
| Cdc14a | 0.31 | 2.29 | -3.089 | 0.004564 | 0.443 |
| Gm13306 | 0.37 | 3.40 | -3.079 | 0.004687 | 0.452 |
| Zfp711 | 0.17 | 4.71 | -3.074 | 0.00474 | 0.453 |
| Mtf2 | 0.14 | 5.98 | -3.073 | 0.004751 | 0.453 |
| Spg20 | -0.17 | 4.45 | 3.068 | 0.004817 | 0.457 |
| Arntl | 0.16 | 4.73 | -3.064 | 0.004865 | 0.458 |
| Shq1 | 0.19 | 2.72 | -3.061 | 0.004902 | 0.459 |
| Tfr2 | 0.33 | 2.25 | -3.057 | 0.004942 | 0.461 |
| Irgm2 | -0.51 | 0.88 | 3.055 | 0.004972 | 0.461 |
| Fgfr3 | -0.16 | 6.38 | 3.052 | 0.005005 | 0.461 |
| Mllt1 | -0.12 | 6.21 | 3.050 | 0.005034 | 0.461 |
| Ccdc14 | 0.28 | 2.15 | -3.048 | 0.005053 | 0.461 |
| Slfn5 | -0.25 | 3.12 | 3.040 | 0.00516 | 0.462 |
| Gm22009 | 0.56 | -0.30 | -3.040 | 0.005165 | 0.462 |
| Tinagl1 | -0.49 | 0.98 | 3.039 | 0.005169 | 0.462 |
| Septin4 | -0.17 | 5.57 | 3.033 | 0.005246 | 0.462 |
| Adss | 0.11 | 6.54 | -3.028 | 0.005314 | 0.462 |
| Specc1 | -0.17 | 6.08 | 3.027 | 0.005325 | 0.462 |
| Ccdc141 | -0.21 | 4.85 | 3.026 | 0.00534 | 0.462 |
| H4c14 | -0.63 | 0.50 | 3.024 | 0.005365 | 0.462 |
| Tipin | 0.16 | 3.76 | -3.023 | 0.005376 | 0.462 |
| Zfp874b | -0.20 | 3.50 | 3.020 | 0.00542 | 0.462 |
| Itpkb | -0.16 | 4.66 | 3.018 | 0.005447 | 0.462 |
| Rbm3 | 0.11 | 6.80 | -3.018 | 0.005449 | 0.462 |
| Gng12 | -0.10 | 6.88 | 3.015 | 0.005481 | 0.462 |
| Rnf2 | 0.12 | 5.48 | -3.015 | 0.005491 | 0.462 |
| Bcan | -0.11 | 8.15 | 3.014 | 0.0055 | 0.462 |
| Sertad3 | -0.34 | 0.87 | 3.014 | 0.005505 | 0.462 |
| Fbxo6 | 0.19 | 3.92 | -3.009 | 0.005568 | 0.465 |
| Aass | -0.38 | 1.35 | 3.007 | 0.005601 | 0.465 |
| Tmco3 | -0.10 | 5.06 | 3.001 | 0.005684 | 0.470 |
| Malat1 | 0.26 | 9.38 | -2.998 | 0.00572 | 0.470 |
| Gnl2 | 0.09 | 5.98 | -2.990 | 0.005842 | 0.472 |
| Hmgb1 | 0.14 | 6.67 | -2.989 | 0.005843 | 0.472 |
| Casp7 | 0.20 | 3.17 | -2.988 | 0.005866 | 0.472 |
| Lamc1 | -0.23 | 6.17 | 2.987 | 0.005881 | 0.472 |
| Irag1 | -0.54 | 0.63 | 2.986 | 0.0059 | 0.472 |
| Hapln4 | -0.30 | 3.94 | 2.983 | 0.005932 | 0.472 |
| 2810402E24Rik | 0.36 | 1.68 | -2.981 | 0.005971 | 0.472 |
| Sdf2l1 | -0.37 | 2.48 | 2.980 | 0.005976 | 0.472 |
| Eny2 | 0.10 | 6.07 | -2.976 | 0.006036 | 0.472 |
| Apln | -0.24 | 4.40 | 2.976 | 0.006038 | 0.472 |
| Bgn | -0.27 | 3.96 | 2.976 | 0.006046 | 0.472 |
| Lonrf3 | -0.35 | 2.86 | 2.968 | 0.006165 | 0.477 |
| Gm45716 | 0.30 | 1.81 | -2.967 | 0.006169 | 0.477 |
| Gab1 | -0.14 | 4.94 | 2.958 | 0.006308 | 0.485 |
| Top2b | 0.10 | 7.63 | -2.952 | 0.006409 | 0.491 |
| Tpp1 | -0.11 | 5.92 | 2.949 | 0.006455 | 0.491 |
| Tagap | 0.42 | 0.02 | -2.946 | 0.006496 | 0.491 |
| Lipa | -0.20 | 4.04 | 2.946 | 0.006497 | 0.491 |
| Rassf10 | -0.30 | 2.19 | 2.941 | 0.006574 | 0.493 |
| She | -0.32 | 2.22 | 2.941 | 0.006585 | 0.493 |
| Tjp2 | -0.13 | 4.93 | 2.937 | 0.006645 | 0.495 |
| Selplg | -0.21 | 4.44 | 2.933 | 0.006709 | 0.495 |
| Tmem64 | -0.13 | 5.54 | 2.933 | 0.006714 | 0.495 |
| Mrc1 | -0.37 | 2.40 | 2.932 | 0.006727 | 0.495 |
| Sgk2 | -0.40 | 0.64 | 2.930 | 0.006752 | 0.495 |
| Wdr5 | 0.09 | 5.53 | -2.927 | 0.006806 | 0.495 |
| Wipf1 | -0.13 | 3.93 | 2.926 | 0.006821 | 0.495 |
| Fbxl5 | 0.11 | 6.69 | -2.925 | 0.00684 | 0.495 |
| Gm10863 | -0.31 | 0.72 | 2.923 | 0.006878 | 0.495 |
| Pou2f2 | 0.17 | 4.35 | -2.916 | 0.006995 | 0.500 |
| Rars | 0.12 | 5.40 | -2.914 | 0.007026 | 0.500 |
| Ighd | -0.57 | -0.06 | 2.914 | 0.007028 | 0.500 |
| Xrra1 | 0.37 | 0.49 | -2.907 | 0.00715 | 0.506 |
| Tal1 | -0.29 | 1.32 | 2.897 | 0.007326 | 0.514 |
| Zfp366 | -0.34 | 3.50 | 2.895 | 0.007361 | 0.514 |
| Cotl1 | 0.14 | 5.91 | -2.893 | 0.007395 | 0.514 |
| Vegfa | -0.13 | 6.28 | 2.891 | 0.007431 | 0.514 |
| Rbms2 | -0.22 | 2.98 | 2.890 | 0.007448 | 0.514 |
| 1810024B03Rik | 0.46 | -0.09 | -2.889 | 0.007467 | 0.514 |
| Pou6f2 | -0.29 | 1.13 | 2.888 | 0.007494 | 0.514 |
| Marchf7 | 0.10 | 6.35 | -2.887 | 0.007505 | 0.514 |
| Dusp5 | 0.33 | 2.66 | -2.883 | 0.007581 | 0.516 |
| Cbr2 | -0.71 | -0.37 | 2.882 | 0.007596 | 0.516 |
| Gm49492 | 0.27 | 3.01 | -2.878 | 0.007666 | 0.516 |
| Unc5b | -0.21 | 4.21 | 2.878 | 0.007666 | 0.516 |
| Pcx | -0.15 | 5.67 | 2.871 | 0.007798 | 0.521 |
| Kcnj10 | -0.19 | 7.61 | 2.870 | 0.007828 | 0.521 |
| Arpc1b | -0.17 | 4.54 | 2.870 | 0.007828 | 0.521 |
| 4833422C13Rik | 0.17 | 3.61 | -2.868 | 0.007855 | 0.521 |
| A230051N06Rik | 0.39 | 0.80 | -2.859 | 0.008039 | 0.528 |
| Gm9403 | -0.83 | -1.21 | 2.855 | 0.008117 | 0.528 |
| Cd93 | -0.30 | 4.84 | 2.853 | 0.00815 | 0.528 |
| Suco | 0.10 | 5.75 | -2.851 | 0.008181 | 0.528 |
| Slc38a2 | -0.13 | 6.63 | 2.851 | 0.008186 | 0.528 |
| Eng | -0.15 | 4.95 | 2.848 | 0.008253 | 0.528 |
| Gm3667 | 0.76 | -1.22 | -2.848 | 0.008257 | 0.528 |
| D330023K18Rik | -0.38 | 0.72 | 2.847 | 0.008268 | 0.528 |
| Ly6c1 | -0.17 | 4.14 | 2.843 | 0.008343 | 0.528 |
| Hic1 | -0.42 | 1.48 | 2.842 | 0.008369 | 0.528 |
| Fchsd1 | 0.25 | 2.85 | -2.842 | 0.008373 | 0.528 |
| Gm50092 | -0.65 | -0.50 | 2.839 | 0.008418 | 0.528 |
| Ddx56 | 0.10 | 4.93 | -2.837 | 0.008465 | 0.528 |
| Niban1 | -0.24 | 2.74 | 2.837 | 0.008473 | 0.528 |
| Cirbp | 0.20 | 6.37 | -2.836 | 0.008484 | 0.528 |
| Sec22a | 0.17 | 4.21 | -2.835 | 0.008515 | 0.528 |
| Eml1 | -0.14 | 5.96 | 2.834 | 0.00853 | 0.528 |
| Cldn20 | 0.75 | -1.00 | -2.834 | 0.00853 | 0.528 |
| Zfx | 0.12 | 5.15 | -2.830 | 0.008608 | 0.530 |
| Msra | 0.18 | 4.52 | -2.829 | 0.008626 | 0.530 |
| Acsbg1 | -0.11 | 6.88 | 2.827 | 0.008673 | 0.530 |
| Zfp276 | -0.17 | 3.93 | 2.827 | 0.008685 | 0.530 |
| Sbsn | 0.46 | 1.58 | -2.822 | 0.008784 | 0.534 |
| Ehd4 | -0.14 | 3.74 | 2.819 | 0.008835 | 0.534 |
| Gm3739 | 0.34 | 1.07 | -2.815 | 0.008928 | 0.534 |
| Eps8 | -0.11 | 5.29 | 2.812 | 0.00899 | 0.534 |
| Lrp4 | -0.29 | 5.42 | 2.812 | 0.008993 | 0.534 |
| Tnfrsf1b | -0.22 | 1.95 | 2.808 | 0.00907 | 0.534 |
| Creb3l2 | -0.21 | 3.87 | 2.808 | 0.00909 | 0.534 |
| Mdp1 | 0.12 | 4.90 | -2.805 | 0.009145 | 0.534 |
| Zc3h6 | 0.22 | 4.24 | -2.804 | 0.009162 | 0.534 |
| Dhx36 | 0.10 | 6.95 | -2.801 | 0.00923 | 0.534 |
| Lrrcc1 | 0.14 | 4.91 | -2.799 | 0.009279 | 0.534 |
| Gm3604 | 0.42 | 0.86 | -2.799 | 0.009286 | 0.534 |
| Ccdc81 | 0.35 | 0.89 | -2.797 | 0.009317 | 0.534 |
| Znhit3 | 0.14 | 3.57 | -2.795 | 0.009358 | 0.534 |
| Myo18b | -0.28 | 1.94 | 2.795 | 0.009358 | 0.534 |
| Gm43844 | 1.31 | -0.79 | -2.795 | 0.009367 | 0.534 |
| C230035I16Rik | 0.59 | -0.62 | -2.794 | 0.009392 | 0.534 |
| Anxa5 | -0.23 | 4.83 | 2.793 | 0.009422 | 0.534 |
| Gipr | 0.26 | 2.55 | -2.792 | 0.009426 | 0.534 |
| 9630001P10Rik | 0.33 | 1.62 | -2.792 | 0.00944 | 0.534 |
| Ift81 | 0.13 | 4.68 | -2.791 | 0.009451 | 0.534 |
| Rgs5 | -0.20 | 7.03 | 2.791 | 0.009452 | 0.534 |
| Gm43437 | -0.33 | 0.92 | 2.786 | 0.009563 | 0.538 |
| Hmcn1 | -0.29 | 3.59 | 2.786 | 0.009583 | 0.538 |
| Sync | -0.73 | -0.59 | 2.778 | 0.009749 | 0.543 |
| Rab7b | -0.30 | 2.11 | 2.778 | 0.009752 | 0.543 |
| Mtmr10 | -0.10 | 4.34 | 2.777 | 0.009786 | 0.544 |
| Gm46123 | -0.69 | -0.72 | 2.774 | 0.009858 | 0.545 |
| Zfp53 | 0.21 | 2.37 | -2.773 | 0.009869 | 0.545 |
| Pgpep1 | -0.15 | 3.92 | 2.771 | 0.009926 | 0.546 |
| Ets1 | -0.21 | 4.91 | 2.768 | 0.009989 | 0.548 |

**Supplementary Table S2**. Significantly enriched GO Biological Process pathways associated with morphine treatment (corresponding to P15 NAc downregulated DEGs).

| **Category** | **Description** | **FDR Value** | **Genes** | **Number of Genes** | **Number of Background Genes** | **P Value** |
| --- | --- | --- | --- | --- | --- | --- |
| GO Biological Process | System development | 0.00022 | Plxnb3, Hapln2, Elovl1, Aspa, Slc45a3, Mal, Bgn, Plp1, Cldn11, Fa2h, Mbp, Ugt8a, Gal3st1, Gjb1, Gpr17, Opalin, Cnp, Gjc2, Lox, Myrf, Mag | 21 | 4350 | 2.05E-07 |
| GO Biological Process | Axon ensheathment | 5.33E-11 | Aspa, Mal, Plp1, Cldn11, Fa2h, Mbp, Ugt8a, Gal3st1, Myrf, Mag | 10 | 116 | 4.14E-15 |
| GO Cellular Component | Myelin sheath | 8.17E-08 | Plp1, Cldn11, Mbp, Mobp, Ermn, Cnp, Mog, Gjc2, Mag | 9 | 212 | 4.82E-11 |
| GO Biological Process | Oligodendrocyte differentiation | 2.69E-07 | Aspa, Plp1, Fa2h, Gpr17, Cnp, Myrf, Mag | 7 | 82 | 8.33E-11 |
| GO Biological Process | Sphingolipid biosynthetic process | 0.00016 | Elovl1, Elovl7, Fa2h, Ugt8a, Gal3st1 | 5 | 71 | 1.27E-07 |
| GO Biological Process | Regulation of gliogenesis | 0.002 | Aspa, Slc45a3, Opalin, Gjc2, Mag | 5 | 147 | 0.00000397 |
| GO Molecular Function | Structural constituent of myelin sheath | 0.0000169 | Mal, Plp1, Mbp, Mobp | 4 | 10 | 5.23E-09 |
| GO Cellular Component | Paranode region of axon | 0.0011 | Ermn, Gjc2, Mag | 3 | 15 | 0.00000293 |
| GO Cellular Component | Internode region of axon | 0.0088 | Mbp, Ermn | 2 | 4 | 0.0000365 |
| GO Biological Process | Central nervous system myelin maintenance | 0.0134 | Fa2h, Myrf | 2 | 4 | 0.0000365 |
| GO Cellular Component | Myelin sheath adaxonal region | 0.0144 | Cnp, Mag | 2 | 6 | 0.0000679 |
| GO Biological Process | Protein localization to paranode region of axon | 0.0178 | Mal, Ugt8a | 2 | 5 | 0.000051 |
